# Topoisomerase-mediated resolution of negative DNA supercoiling sustains human transcription and gene activation

**DOI:** 10.64898/2026.09.30.755832

**Authors:** Jun Lyu, Abdelmonsif Ahmed, Ipsita Biswas, Shuai Li, Chongyi Chen

## Abstract

Transcription generates DNA supercoiling that can influence gene expression, but how transcription-associated supercoiling shapes RNA polymerase II (RNAPII) dynamics in human cells remain poorly understood. Here, we show that topoisomerase-mediated resolution of negative DNA supercoiling is required for productive transcription. Acute topoisomerase perturbation causes negative supercoiling to accumulate around transcription start sites (TSS), with greater negative supercoiling accumulation associated with stronger transcription reduction. Elevated negative supercoiling behind the elongating RNAPII, but not positive supercoiling ahead, is associated with reduced RNAPII elongation rates. Using NOTE-seq2 to simultaneously profile newly synthesized and total RNA in single cells, we further demonstrate that acute TOP1 depletion increases gene expression noise by reducing both transcriptional burst size and frequency. Mechanistically, negative supercoiling accumulation at TSS increases RNAPII promoter occupancy and promoter-proximal pausing, providing a potential basis for reduced burst size. Micro-C analysis shows that TOP1 depletion largely preserves chromatin compartments and TADs but weakens short-range chromatin loops, including enhancer-promoter interactions, likely contributing to reduced burst frequency. Finally, during T-cell activation, impaired resolution of negative supercoiling compromises rapid gene upregulation and attenuates the transcriptional activation program. Together, these findings establish topoisomerase-mediated resolution of negative DNA supercoiling as a key mechanism linking DNA topological homeostasis to RNAPII dynamics, chromatin looping, and rapid gene activation.

## Introduction

Transcription requires RNA polymerase to translocate along the helical DNA template and therefore inevitably generates torsional stress. As RNA polymerase progresses, positive supercoiling accumulates ahead of the transcription complex, whereas negative supercoiling is generated behind it, as described by the twin-supercoiled-domain model^1,2^. In turn, accumulated DNA supercoiling around genes, if not efficiently resolved, can profoundly influence gene expression. Negative supercoiling has been shown to generally facilitate transcription by promoting DNA melting at promoters and preinitiation complex assembly, whereas positive supercoiling can impede transcriptional elongation by generating torsional resistance^3–12^. However, this general understanding of supercoiling’s impact on transcription has been derived primarily from in vitro systems^3,4^, microorganisms^8–11^, or exogenous reporter constructs of specific genes^7,12^. The genome-wide principles underlying endogenous human genes’ response to DNA supercoiling accumulation remain unclear.

DNA supercoiling is tightly regulated by topoisomerases^13,14^, and supercoiling alterations upon topoisomerase perturbation have been used to investigate the effect of DNA supercoiling on transcription. However, previous studies often relied on prolonged depletion or topoisomerase poisons^15–23^, which can introduce secondary effects, including chromatin adaptation, replication stress, and DNA damage, complicating the attribution of observed transcriptional changes directly to altered DNA supercoiling^24–26^. Recent studies using acute perturbations have begun to address these limitations. Acute depletion of topoisomerases in C. elegans using degron systems has revealed widespread transcriptional defects following disruption of DNA topology^27^. Studies in yeast have further implicated negative DNA supercoiling in transcriptional bursting and in coupling the transcriptional behavior of neighboring genes^23^. Our work found that negative supercoiling accumulated around human genes is associated with transcriptional repression^28^, rather than facilitating transcription. However, technical features of the experimental system, including basal degron leakage before induction and changes associated with cell-cell compositions upon acute topoisomerase depletion, particularly involving cells undergoing DNA replication or mitotic chromosome condensation, could complicate the interpretation. Thus, it remains to be determined how disrupted DNA topological homeostasis in human cells influences endogenous transcriptional activity, and which steps of the transcription process are particularly sensitive to DNA supercoiling alterations.

In this work, we perturbed DNA supercoiling by acutely depleting TOP1 in G1-synchronized human cells and investigated its role in transcription by integrating genome-wide measurements of DNA supercoiling, nascent transcription activity, single-cell transcription activity and transcriptional bursting, RNAPII occupancy and elongation, and 3D chromatin architecture. We found that accumulated negative DNA supercoiling at the transcriptional start sites (TSS) is associated with human transcription repression. Negative, rather than positive, supercoiling is strongly associated with reduced RNAPII elongation and increased promoter-proximal pausing. We further developed NOTE-seq2 to resolve single-cell transcriptional dynamics and found that TOP1 depletion reduces transcriptional burst frequency and burst size. Micro-C results reveal weakened local chromatin loops and compromised enhancer-promoter interactions. Finally, we extend these findings to the process of T-cell activation, showing that genes undergoing rapid transcriptional activation are particularly sensitive to acute TOP1 depletion. Together, our results highlight the active maintenance of DNA topological homeostasis, particularly the efficient resolution of accumulated negative supercoiling, is crucial to sustain human transcription and rapid gene activation.

## Results

### Negative supercoiling accumulation upon topoisomerase perturbation represses human transcription

To investigate the impact of DNA supercoiling on transcription, we acutely depleted TOP1 in HCT116 cells using a TOP1 degron system to globally increase DNA supercoiling and examined the resulting changes in transcriptional activity (Fig. 1A, B)^26,29^. Importantly, because DNA replication can also contribute to and be affected by DNA supercoiling as a potential confounding factor, we synchronized cells in G1 phase by lovastatin treatment ahead of supercoiling perturbation^30^. Cell-cycle profiling confirmed efficient G1 synchronization, and TOP1 was acutely depleted in the synchronized cells following two hours of auxin (IAA) treatment (Supplementary Fig. 1). To detect the potentially rapid transcriptional change, we performed TTchem-seq to capture the transcriptional activity immediately following a supercoiling alteration.

**Figure 1.**
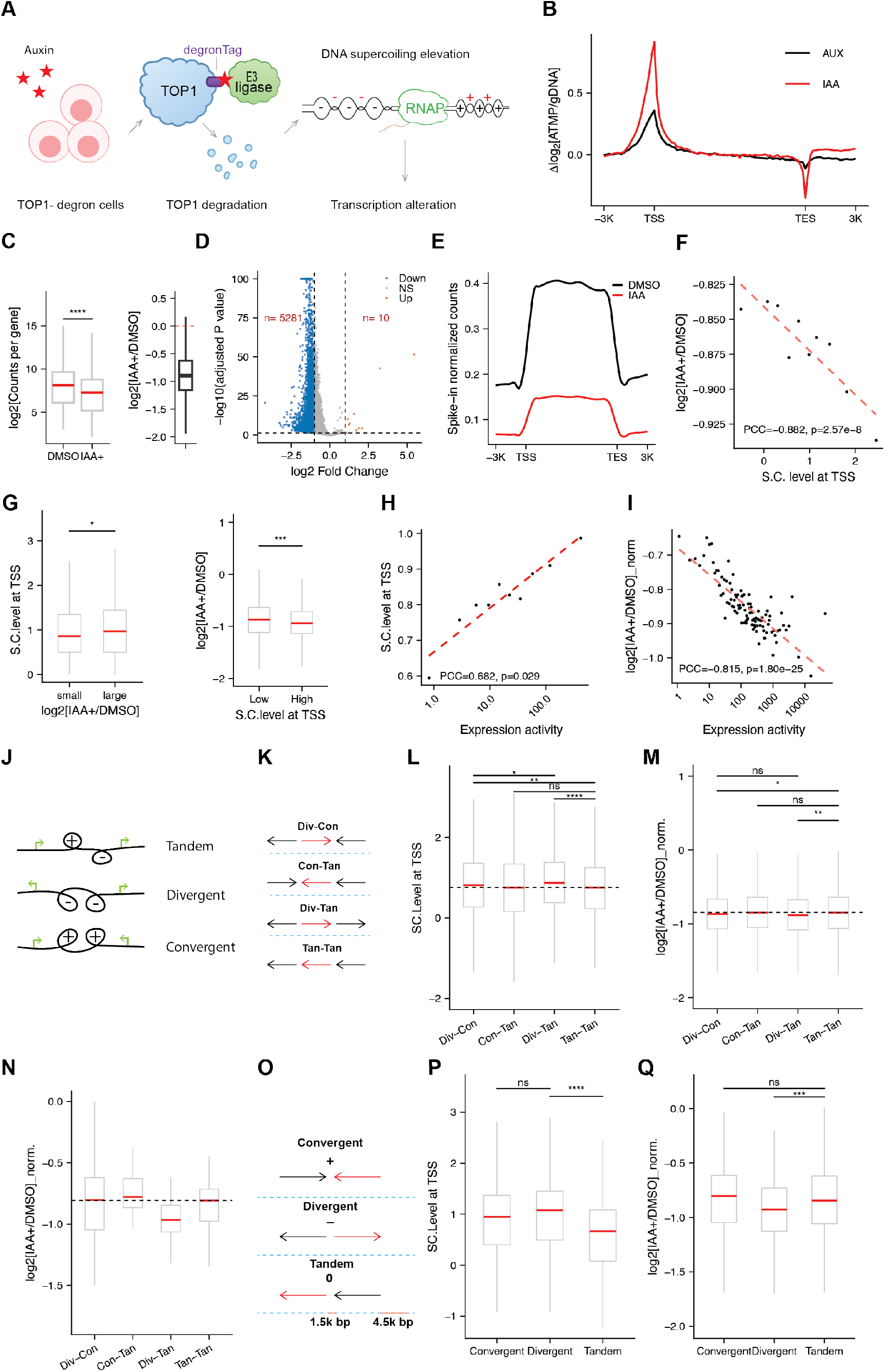
Accumulated negative supercoiling upon TOP1 depletion represses transcription. **A.** Schematic of the TOP1 degron system in the HCT116 cell line. **B.** Supercoiling level around genes upon IAA (TOP1 depletion) or AUX (Auxinole, control) treatment, relative to the supercoiling level at the gene body. **C.** Left: Transcription activity of genes upon TOP1 depletion (IAA) versus control (DMSO), measured by TTchem-seq. Right: Fold change in transcription activity of genes upon TOP1 depletion. n= 17227 genes. Transcription activity of genes is the average of two replicates. **D.** Genes with significantly changed transcription activity (p-adj ≤ 0.05, |log2(fold change)| ≥ 1, n= 17227 genes). **E.** Transcription activity along the gene body of genes upon TOP1 depletion versus control (n = 17227 genes). **F.** The correlation between supercoiling (S.C.) level at transcription start sites (TSS) and the fold change in transcriptional activity following TOP1 depletion relative to the DMSO control. Genes were grouped into bins of 1,000 genes, with each dot representing one bin. The dashed red line indicates the linear regression fit (n= 10 bins for 9483 genes). **G.** Left: Comparison of S.C. level in DMSO control at the TSS between genes showing the smallest and largest changes in transcriptional activity upon TOP1 depletion relative to DMSO control. Right: Comparison of fold change in transcriptional activity between genes with the lowest and highest supercoiling levels in DMSO control. Each category includes 1000 genes. Red lines indicate medians. **H.** The correlation between S.C. level at the TSS and transcriptional activity of genes in the DMSO condition. Genes were grouped into bins of 1,000 genes, with each dot representing one bin. The dashed red line indicates the linear regression fit (n= 10 bins for 9483 genes). **I.** The correlation between basal transcriptional activity of genes in DMSO condition and the fold change in transcriptional activity of genes upon TOP1 depletion (IAA). The dashed red line indicates the linear regression fit. Genes were grouped into bins of 100 genes, with each dot representing one bin (n=102 bins for 10121 genes). **J.** Schematic of transcription-induced supercoiling at different gene-pair orientations. Arrows indicate transcription direction, while “+” and “−” denote positive and negative supercoiling, respectively. **K.** Schematic of gene context based on the transcriptional orientation of neighboring genes. Black and red arrows indicate the transcriptional directions of the neighboring and focal genes, respectively. **L and P.** S.C. level at the TSS of genes across different gene contexts. **M, N, and Q.** The normalized fold change in transcriptional activity upon TOP1 depletion (IAA) across different gene contexts. The distance between the neighboring and focal genes is less than 1,500 bp in (N), whereas in (Q), the distance is greater than 4,500 bp on one side and less than 1,500 bp on the other. In (L), (M), (N), and (Q), red horizontal lines indicate group medians, and the dashed line indicates the median of the group in the tandem context. **O.** Schematic of gene context based on the transcriptional orientation of neighboring genes within 1,500 bp. Arrows indicate transcription direction, while “+” and “−” denote positive and negative supercoiling, respectively. The “0” denotes a relatively neutral supercoiling state. Distances of 1.5 kb and 4.5 kb indicate the regions considered relative to the neighboring genes. In (L) and (M), n= 2774 genes for Div-Con; n=2406 genes for Con-Tan; n=2413 genes for Div-Tan; n=2466 genes for Tan-Tan. In (N), n= 46 genes for Div-Con; n=24 genes for Con-Tan; n=31 genes for Div-Tan; n=31 genes for Tan-Tan. In (P) and (Q), n= 316 genes for Convergent; n= 579 genes for Divergent; n= 404 genes for Tandem. Pearson correlation coefficient (PCC) and associated *P* value are shown as indicated in (F), (H), and (I). Statistical significance is indicated in (C), (G), (H), (L), (M), (P), and (Q) (*P < 0.05, **P < 0.01, ***P < 0.001,****P < 0.0001; ns, not significant).

TTchem-seq revealed a global reduction in transcriptional activity after TOP1 depletion (Fig. 1C-E), with 5,281 genes significantly downregulated compared with only 10 genes upregulated under the indicated thresholds (Fig. 1D). Such a global transcriptional reduction was further confirmed by an orthogonal approach using SLAM-seq, measuring the newly synthesized mature RNA (Supplementary Fig. 2A). Moreover, the reduction of nascent transcripts was evenly distributed across the gene body from TSS to TES, suggesting a predominant transcriptional defect starting from the 5’ end of the genes (Fig. 1E). We therefore reasoned that the transcription reduction is associated with supercoiling at the TSS, where negative DNA supercoiling dramatically accumulates upon TOP1 depletion (Fig. 1B), instead of the roles of topoisomerase and supercoiling in RNAP elongation. Indeed, we found that the fold change in transcriptional activity following TOP1 depletion was strongly correlated with the level of negative supercoiling at the TSS, with genes exhibiting a greater transcriptional reduction showing higher TSS supercoiling levels than those exhibiting a smaller reduction, and genes with higher TSS supercoiling levels showing a greater reduction in transcriptional activity than those with lower TSS supercoiling levels (Fig. 1F, G). In addition, transcriptional activity was positively correlated with TSS supercoiling levels, with highly transcribed genes exhibiting higher levels of supercoiling at the TSS, consistent with our previous findings and the twin-supercoiled-domain model (Fig. 1H)^1,29^. We then used transcriptional activity as a proxy for TSS supercoiling levels to stratify genes by TSS supercoiling levels at higher resolution. Similar to the correlation between directly observed TSS supercoiling levels and changes in transcriptional activity observed at the 1,000-gene scale (Fig. 1F), inferred TSS supercoiling levels from transcriptional activity were strongly correlated with changes in transcriptional activity at the 100-gene scale, with highly transcribed genes exhibiting a greater reduction in transcriptional activity following TOP1 depletion (Fig. 1I).

DNA supercoiling can propagate along chromatin and thereby influence the topological state of neighboring genes, creating distinct supercoiling environments depending on neighboring-gene orientation (Fig. 1J, K)^5,10,31–33^. We therefore used gene context as an additional proxy for TSS supercoiling levels, allowing us to examine the relationship between supercoiling and transcription while minimizing potential confounding effects of the noncanonical functions of TOP1. Based on the relative orientations of the flanking genes and the focal gene, we classified focal genes into four gene-context categories: divergent–convergent, convergent–tandem, divergent–tandem, and tandem–tandem. We matched transcriptional activity across these four categories to minimize the contribution of transcriptional activity itself and thereby better isolate the effect of gene context (Supplementary Fig. 2B). We reasoned that focal genes involving a divergent configuration would exhibit higher levels of supercoiling at the TSS than those involving a tandem configuration. Consistent with this prediction, genes in the divergent– convergent and divergent–tandem contexts exhibited higher TSS supercoiling levels and, correspondingly, a greater reduction in transcriptional activity following TOP1 depletion (Fig. 1L, M). To enhance the effect of gene context arising from supercoiling propagation, we restricted the neighboring genes to within 1.5 kb. We found genes in the divergent-tandem context exhibited more obvious reduction than genes in other gene contexts upon TOP1 depletion (Fig. 1N and Supplementary Fig. 2C). Moreover, we restricted one neighboring gene to within 1.5 kb of the focal gene and the other to more than 4.5 kb away, thereby strengthening the supercoiling contribution from the proximal neighbor while minimizing that from the distal neighbor (Fig. 1O). We classified genes into three categories with matched transcriptional activity (Supplementary Fig. 2D): convergent, divergent, and tandem. Consistent with the previous prediction, genes in a divergent context exhibit a higher supercoiling level and, correspondingly, a greater reduction in transcriptional activity following TOP1 depletion (Fig. 1P, Q).

To further validate the relationship between DNA supercoiling and transcription, we asked whether this phenomenon could be reproduced using an orthogonal approach for supercoiling perturbation. We treated G1-synchronized HCT116 cells with the TOP2 inhibitor ICRF193, which resulted in an increase in gene-associated supercoiling accompanied by a global reduction in nascent transcription, recapitulating the phenomenon following acute TOP1 depletion (Supplementary Fig. 3A–D). In addition, basal TSS supercoiling was associated with transcriptional activity, and highly transcribed genes again exhibited greater reductions in transcriptional activity following TOP2 inhibition (Supplementary Fig. 3E, F). Importantly, analysis using gene context to infer TSS supercoiling also recapitulated the relationship between supercoiling levels and transcriptional responses following TOP2 perturbation (Supplementary Fig. 3G–I). Together, these results demonstrate a robust association between negative supercoiling accumulation at the TSS and reduced transcriptional activity, highlighting an inhibitory role of negative supercoiling on transcription in human cells.

### Negative supercoiling behind RNAPII, but not positive supercoiling ahead, slows transcription elongation

Positive supercoiling generated ahead of an elongating RNA polymerase has long been recognized as a topological barrier to transcriptional elongation^7–9,18,33^. Because acute TOP1 depletion increased both negative supercoiling at the TSS and positive supercoiling at the TES (Fig. 1B), we asked whether the accumulation of positive supercoiling at the TES influences transcription elongation following TOP1 depletion.

To assess its impact on transcription elongation, we profiled Ser2-phosphorylated RNAPII (RNAPII-Ser2P), a marker of elongating RNAPII. Not surprisingly, despite the global reduction in nascent transcription, TOP1 depletion increased RNAPII-Ser2P occupancy across transcribed regions, with particularly prominent accumulation toward the TES, suggesting impaired polymerase progression and slower transcription elongation (Fig. 2A and Supplementary Fig. 4A). To quantify changes in transcriptional elongation, we developed the elongation index based on the ratio of RNAPII-Ser2P occupancy to nascent RNA production (Fig. 2B). The elongation index reflects the amount of transcriptionally engaged RNAPII required to produce a given amount of nascent RNA, with a higher elongation index indicating slower RNAPII progression and a lower elongation index indicating faster progression (Fig. 2B). Accordingly, the accumulation of RNAPII-Ser2P without a proportional increase in nascent RNA production following acute TOP1 depletion resulted in an increased elongation index, suggesting a slower RNAPII progression (Fig. 2B, C).

**Figure 2.**
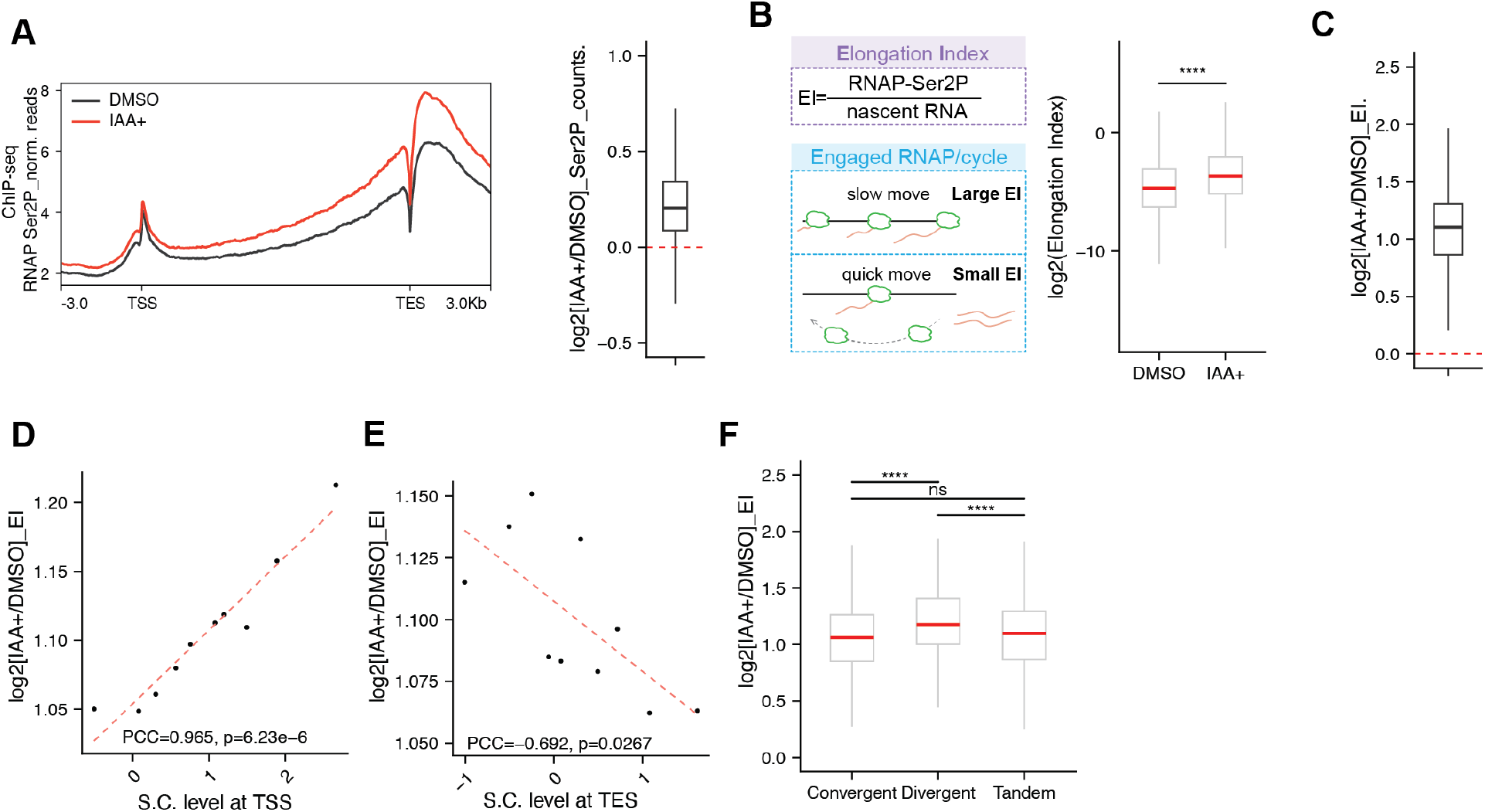
Negative supercoiling behind RNAPII, but not positive supercoiling ahead, slows transcription elongation. **A.** Left: Normalized RNAPII-Ser2P ChIP-seq signal at genes in DMSO control and TOP1 depletion cells. Right: Fold changes in Ser2P signal of genes upon TOP1 depletion (n= 9916 genes). The ChIP-seq signal at genes is the average of two replicates. **B.** Left: Schematic of the elongation index describing the speed of elongating RNAPII. Right: elongation indices of genes in control and TOP1-depleted cells (n= 9916 genes). **C.** Fold change in elongation indices of genes upon TOP1 depletion (n= 9916 genes). **D-E.** The correlation between S.C. level at TSS (D) or TES (E) and the fold change in elongation indices of genes upon TOP1 depletion relative to control. Genes were grouped into bins of 1,000 genes, with each dot representing one bin. The dashed red line indicates the linear regression fit (n= 10 bins for 9916 genes). **F.** The fold change in elongation indices of genes upon TOP1 depletion across different gene contexts (n= 295 genes for Convergent; n= 453 genes for Divergent; n= 391 genes for Tandem). In (B) and (F), red horizontal lines indicate group medians, with statistical significance indicated (****P < 0.0001; ns, not significant). Pearson correlation coefficient (PCC) and associated *P* value are shown as indicated in (D) and (E).

Interestingly, the change in elongation index following TOP1 depletion was strongly associated with supercoiling level at the TSS, whereas supercoiling level at the TES showed a weaker, less-significant relationship (Fig. 2D, E). Reciprocal stratification analyses further supported the relationship between the change in elongation index and TSS supercoiling, whereas no significant association was observed with TES supercoiling (Supplementary Fig. 4B, C). We also classified genes with matched transcriptional activity into three gene contexts with defined distance constraints (Fig. 1O and Supplementary Fig. 4D). In this framework, the convergent context enhances positive supercoiling at the TES, whereas the divergent context enhances negative supercoiling at the TSS, allowing gene context to serve as a proxy for the relative contributions of positive and negative supercoiling. Gene context-based analysis revealed that genes in the divergent context exhibited a greater increase in the elongation index following TOP1 depletion than genes in the convergent context, whereas the change in elongation index was comparable between genes in the convergent and tandem contexts (Fig. 2F). Thus, enhanced negative supercoiling associated with the divergent context was linked to impaired RNAPII progression. Together, these results indicate that the impaired RNAPII elongation following TOP1 depletion is primarily associated with negative supercoiling accumulation behind the elongating RNAPII at the TSS rather than positive supercoiling accumulation ahead at the TES.

### NOTE-seq2 reveals that acute TOP1 depletion increases gene expression noise and reduces transcriptional burst size and burst frequency

Having established that negative supercoiling is associated with impaired transcriptional activity, we next sought to determine how this topological constraint affects transcriptional dynamics at the single-cell level. Previous studies have implicated DNA supercoiling in regulating transcriptional bursting at individual genes in microorganisms^9,23^. However, whether DNA supercoiling globally regulates transcriptional bursting in human cells remains unclear. We developed NOTE-seq2, an improved version of NOTE-seq that enables simultaneous profiling of newly synthesized and total RNA in individual cells (Fig. 3A)^34^. NOTE-seq2 incorporates a cell-sorting step, enabling enrichment of the cell population of interest while removing cell debris and multiplets (Fig. 3A). In addition, NOTE-seq2 is compatible with the latest 10x Genomics 3′ Gene Expression platform. Together, these improvements reduced the multiplet rate from 10.2% to 1.1% and approximately doubled the number of transcripts detected per cell compared with the original NOTE-seq method (Fig. 3B–E).

**Figure 3.**
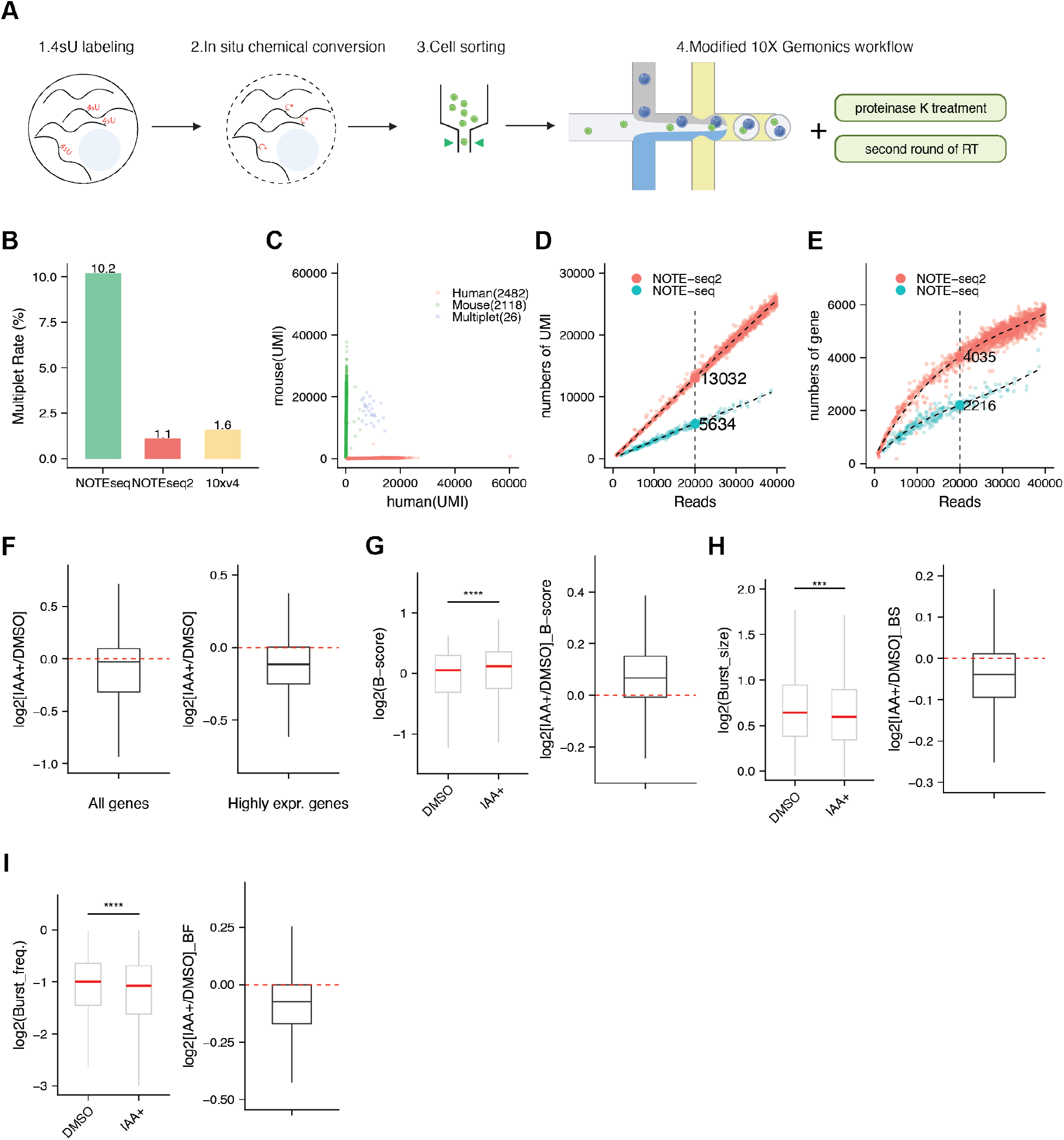
Acute TOP1 depletion elevates gene expression noise and reduces transcriptional burst size and burst frequency. **A.** Schematic of NOTE-seq2. **B.** Comparison of the multiplet rates between NOTE-seq, NOTE-seq2, and 10xv4 3’ expression kit. **C.** Transcript counts of individual cells mapped to the merged human-mouse genome, with cells colored based on their assigned categories of human, mouse, or mix. Cell numbers are indicated. **D-E.** Comparison of detected UMI (D) or gene (E) numbers per cell (NIH3T3) between NOTE-seq and NOTE-seq2. **F.** Fold changes of newly synthesized RNA upon TOP1 depletion in G1-sorted cells. **G-I.** Changes in transcriptional bursting following TOP1 depletion. (G) Expression noise (B-score) in DMSO- and IAA-treated cells and the distribution of log2 fold changes in burst score following TOP1 depletion relative to DMSO. (H) Burst size and the corresponding log2 fold changes following TOP1 depletion. (I) Burst frequency (B-score) and the corresponding log2 fold changes following TOP1 depletion. N= 2120 genes for the top 10% highly expressed genes. Red lines indicate group means, and red dashed lines indicate no change (log2 fold change = 0). Statistical significance is indicated (***P < 0.001, ****P < 0.0001; ns, not significant).

We applied NOTE-seq2 to G1-sorted control and TOP1-depleted cells and detected substantial numbers of genes and RNA molecules in both the total and newly synthesized RNA fractions (Supplementary Fig. 5). Notably, gene and RNA molecule detection in the total RNA fraction was comparable between the two conditions, arguing against systematic differences in transcript detection as an explanation for the changes observed in newly synthesized RNA (Supplementary Fig. 5). Pseudo-bulk analysis of newly synthesized RNA revealed a global reduction following TOP1 depletion, with a more pronounced decrease among highly expressed genes, consistent with our population-level nascent-transcription measurements (Fig. 3F).

We then quantified gene expression noise, transcriptional burst frequency (fraction of transcriptionally active cells) and burst size (mean nascent RNA output per transcriptionally active cell) using newly synthesized RNA, focusing on highly expressed genes to obtain robust estimates of transcriptional dynamics. Acute TOP1 depletion significantly increased gene expression noise (Fig. 3G), while reduced both transcriptional burst size and burst frequency (Fig. 3H, I). Together, these findings link the accumulation of transcription-associated supercoiling upon acute TOP1 depletion to altered transcriptional bursting dynamics.

### Accumulated negative DNA supercoiling increases RNAPII promoter occupancy and promoter-proximal pausing

The reduction in transcriptional burst size following acute TOP1 depletion suggested that, once a transcriptional burst is initiated, fewer RNA molecules is produced during each period of the ON state. We first examined RNAPII occupancy at promoters, where transcription initiation occurs and the transcriptional defect originates (Fig. 1E). RNAPII ChIP-seq revealed pronounced accumulation of RNAPII around the TSS following acute TOP1 depletion (Fig. 4A and Supplementary Fig. 6A). The increase in RNAPII occupancy at the TSS was positively associated with the negative supercoiling level at the TSS (Fig. 4B and Supplementary Fig. 6B, C), suggesting enhanced RNAPII occupancy at promoters upon elevated supercoiling level during acute TOP1 depletion. Moreover, the increase in RNAPII occupancy at the TSS was correlated with the fold change in transcriptional activity following acute TOP1 depletion (Fig. 4C and Supplementary Fig. 6D), suggesting that higher RNAPII occupancy at the TSS indeed stimulates transcription, resulting in less severe transcriptional repression. This observation is consistent with the long-standing concept that negative supercoiling at the TSS facilitates RNAPII recruitment and transcription initiation^3,4^. However, despite increased promoter-associated RNAPII occupancy, overall transcriptional output was reduced following acute TOP1 depletion, raising the possibility that promoter-bound RNAPII is in a promoter-proximal paused state that cannot efficiently transit from initiation into elongation. We therefore calculated the RNAPII pausing index to determine whether acute TOP1 depletion impairs the transition from promoter-proximal pausing to productive transcription. Because the global increase in RNAPII-Ser2P occupancy following TOP1 depletion could confound conventional estimation of the pausing index, we defined an adjusted pausing index as the ratio of promoter-proximal to gene-body RNAPII occupancy normalized by the elongation index (Fig. 4D). Indeed, we found that acute TOP1 depletion significantly increased the adjusted pausing index (Fig. 4E). Importantly, TSS supercoiling level was strongly associated with changes in the adjusted pausing index (Fig. 4F and Supplementary Fig. 6E, F), and greater increases in pausing resulted in more significant reductions in transcriptional activity (Fig. 4G and Supplementary Fig. 6G). Together, these findings support a model in which negative supercoiling accumulation exerts opposing effects on distinct steps of the transcription cycle: it facilitates RNAPII occupancy at the TSS but also increases its promoter-proximal pausing, thereby limiting the transition from transcription initiation to productive elongation and ultimately reducing the overall transcriptional output.

**Figure 4.**
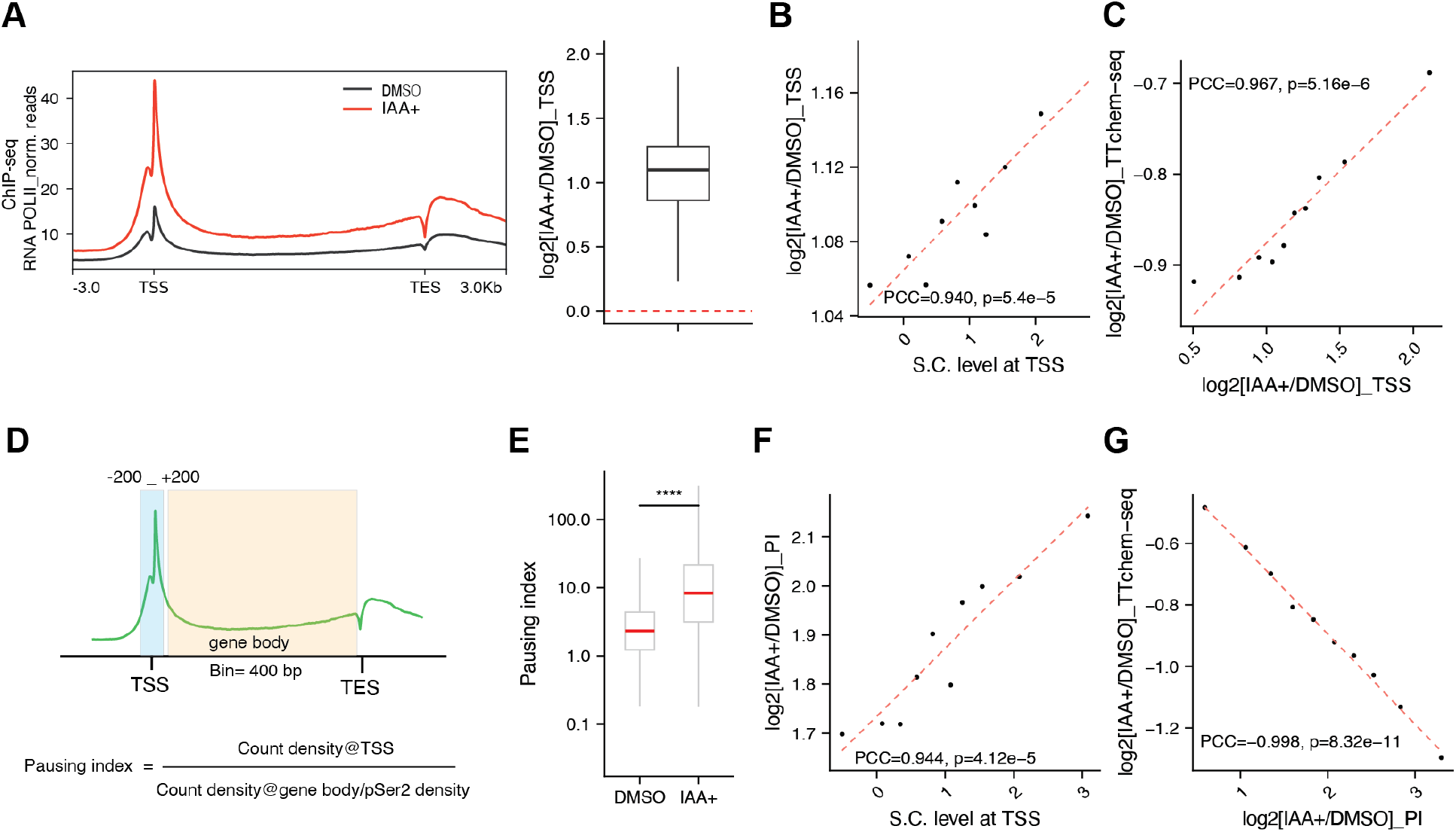
Accumulated negative DNA supercoiling at the TSS increases RNAPII promoter occupancy and promoter-proximal pausing. **A.** Left: Normalized RNAPII ChIP-seq signal at genes in DMSO control and TOP1 depletion cells. Right: Fold changes in RNAPII signal at genes upon TOP1 depletion (n= 9676 genes). The ChIP-seq signal at genes is the average of two replicates. **B-C.** Correlation between S.C. level at the TSS (B, n=10 bins for 9123 genes) or fold change in transcription activity (C, n=10 bins for 9057 genes) and the fold change in RNAPII binding at the TSS of genes upon TOP1 depletion. **D.** Schematic of pausing index calculation. **E.** Pausing indices of genes in DMSO- and IAA-treated cells (n= 9674 genes, ****P < 0.0001). **F-G.** Correlation between S.C. level at the TSS (F, n=10 bins for 9122 genes) or fold change in transcription activity (G, n=10 bins for 9388 genes) and the fold change in pausing indices at the TSS upon TOP1 depletion. In B, C, F, and G, genes were grouped into bins of 1,000 genes, with each dot representing the median of one bin. The dashed red line indicates the linear regression fit. Pearson correlation coefficient (PCC) and associated *P* value are shown as indicated.

### Acute TOP1 depletion compromises chromatin loop formation and enhancer-promoter interactions

The reduction in transcriptional burst frequency following acute TOP1 depletion suggested that DNA supercoiling may also affect upstream OFF-to-ON regulatory processes that influence transcriptional activation. Previous studies have implicated nucleosome occupancy, particularly nucleosomes around the TSS, may be an important regulator of burst frequency^35^. We therefore examined nucleosome occupancy around the TSS using micrococcal nuclease sequencing (MNase-seq). However, acute TOP1 depletion did not result in any obvious change in TSS-associated nucleosome occupancy (Supplementary Fig. 7A), suggesting that altered nucleosome organization is unlikely to account for the observed reduction in transcriptional burst frequency. We next considered whether acute TOP1 depletion affects 3D chromatin architecture. Given the established role of enhancer–promoter interactions in transcriptional bursting, we performed Micro-C to determine whether accumulated supercoiling tension affects chromatin structure and DNA looping.

Via Micro-C analysis, we found that TOP1 depletion leads to no major changes in large-scale chromatin organization, including A/B compartments and TADs (Supplementary Fig. 7B-D), in agreement with our previous report^29^. However, we observed a substantial reduction in small-scale chromatin looping accompanied by an overall decrease in the loop strength upon acute TOP1 depletion (Fig. 5A, B). Notably, the reduction in chromatin loops was comparable between large-scale negative- and positive-supercoiling domains across the human genome, suggesting that the observed loop disruption was not preferentially associated with either supercoiling state of the chromatin (Supplementary Fig. 7E). Importantly, the majority of lost and shrunk chromatin loops involved enhancer and/or promoter regions, suggesting that compromised small-scale chromatin looping impair enhancer–promoter interactions (EPIs) (Fig. 5C). Given the central role of cohesin-mediated loop extrusion in shaping chromatin loops, we next examined chromatin stripes, which reflect extended patterns of chromatin interactions associated with loop-extrusion dynamics. Interestingly, acute TOP1 depletion reduced the number of chromatin stripes, weakened overall stripe strength, and shortened stripe length globally, supporting a model in which compromised topoisomerase activity leads to less processive cohesin-mediated loop extrusion, impairing small-scale chromatin looping including EPIs (Fig. 5D-F).

**Figure 5.**
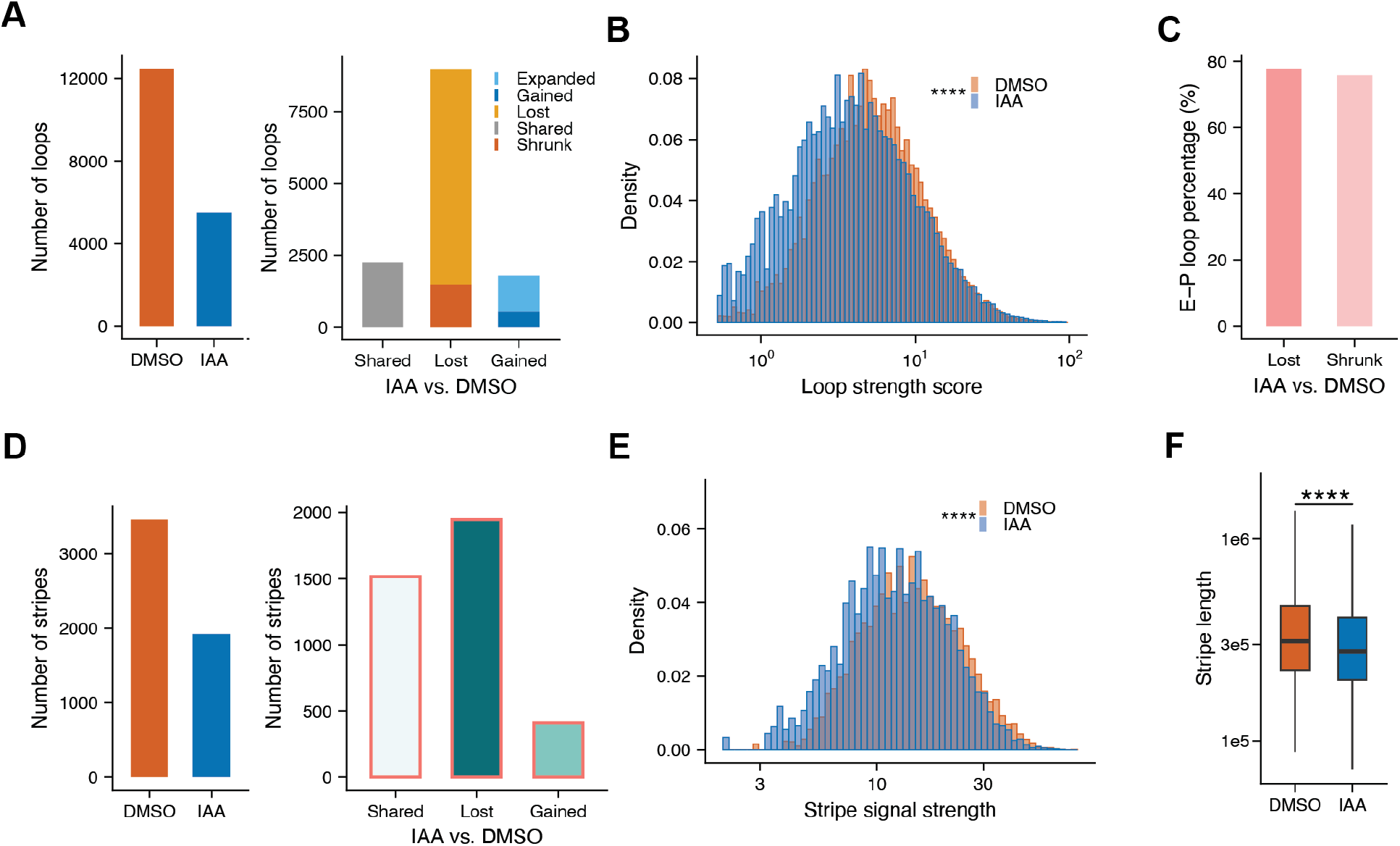
Acute TOP1 depletion compromises chromatin loop formation and enhancer-promoter interactions. **A.** Left: number of chromatin loops identified in DMSO- and IAA-treated cells. Right: number of loop changes upon TOP1 depletion. **B.** The distribution of loop strength scores. **C.** Percentage of enhancer–promoter (E–P) loops among lost and shrunk loops upon TOP1 depletion. **D.** Left: number of chromatin stripes identified in DMSO- and IAA-treated cells. Right: number of stripe changes upon TOP1 depletion. **E.** The distribution of stripe signal strength. **F.** The distribution of stripe lengths in DMSO- and IAA-treated cells (n = 3463 stripes for DMSO and n=2228 stripes for IAA, ****P < 0.0001).

Together, these findings indicate that acute TOP1 depletion preferentially compromises local chromatin 3D structures, including enhancer-promoter contacts, while largely preserving nucleosome occupancy and large-scale genome architecture. The disruption of EPIs provides a potential mechanism contributing to the reduced transcriptional burst frequency.

### Unresolved negative supercoiling compromises rapid gene upregulation upon T-cell activation

Having established that negative supercoiling accumulation represses transcriptional output and alters RNAPII dynamics under steady-state conditions, we next examined whether this topological constraint of the chromatin also affects physiologically relevant gene regulation processes. We used T-cell activation as the model system, which requires drastic activation of stimulus-responsive genes within a short time frame and therefore provides an ideal system to investigate how DNA supercoiling accumulation due to compromised topoisomerase resolution would influence rapid gene activation.

We generated a Jurkat T-cell line with its endogenous TOP1 tagged by an FKBP12^F36V^ degron and mClover, enabling acute TOP1 degradation following dTAGv1 treatment (Fig. 6A-C and Supplementary Fig. 8A). We first examined the consequences of acute TOP1 depletion under steady-state conditions. Consistent with our findings in HCT116 cells, TOP1 depletion increased transcription-associated DNA supercoiling level in Jurkat cells (Supplementary Fig. 8B), as well as a global reduction in nascent transcription measured by SLAM-seq (Supplementary Fig. 8C, D). Thus, Jurkat cells recapitulated the phenomenon in HCT116 cells that elevated negative supercoiling represses transcriptional output. We next examined whether and how this topological constraint may affect rapid gene activation. We activated Jurkat cells with PMA and ionomycin for one hour, producing a rapid transcriptional response, with 541 genes upregulated and 223 genes downregulated (Fig. 6D). To distinguish the effects of supercoiling on dynamically activated versus steady-state genes, we defined significantly upregulated genes as responsive genes (RSGs) and selected a stringent set of unchanged house-keeping genes as non-responsive genes (NRSGs) for subsequent analyses.

**Figure 6.**
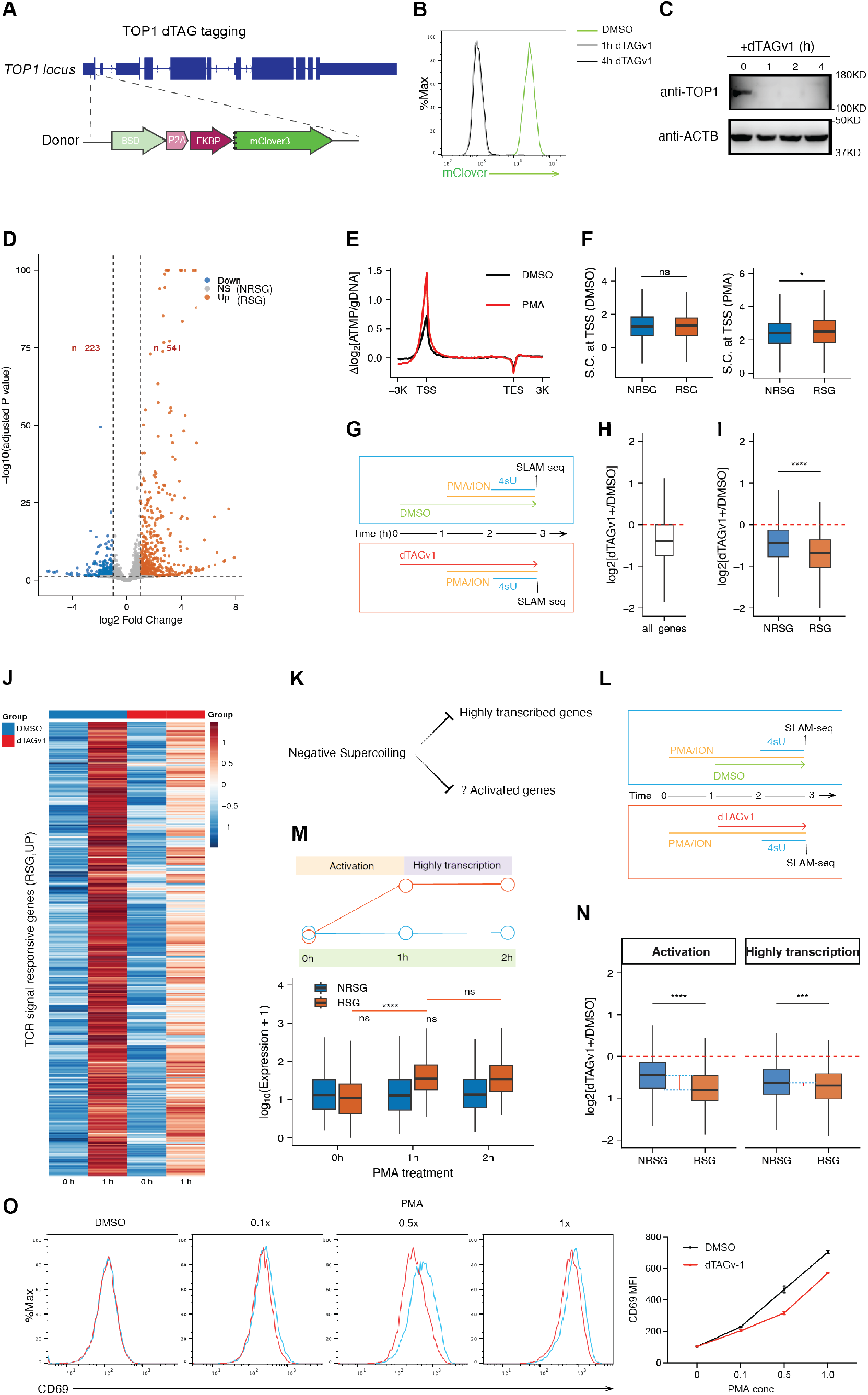
Unresolved negative supercoiling compromises rapid gene upregulation upon T cell activation. **A.** Schematic of dTAG tagging for TOP1. **B-C.** Efficient depletion of mClover-tagged TOP1 following 1- or 4-hour dTAGv1 treatment, measured by flow cytometry (B) and western blot (C). **D.** Differential gene expression upon 1-hour T cell activation by PMA and ionomycin (hereafter, PMA). Significantly upregulated genes and downregulated genes are shown in orange and blue, respectively, while non-significantly regulated genes are shown in gray. Dashed lines indicate the thresholds used to define differential expression. NRSG (n= 3779 genes) is a subset of NS genes defined using the indicated criteria (padj > 0.1 and |log2FoldChange| ≤ 0.25). RSG (n= 541 genes) includes all significantly upregulated genes. The analysis was based on three replicates. **E.** Supercoiling level around genes upon T cell activation (PMA) or control (DMSO), relative to the supercoiling level at the gene body. **F.** S.C. levels at TSSs of non-responsive genes (NRSG) and responsive genes (RSG) under DMSO and PMA conditions. Statistical significance between groups is indicated above the boxplots (*P* < 0.05; ns, not significant). **G.** Schematic of the experimental workflow. Control cells were treated with DMSO, whereas TOP1 depletion was induced with dTAGv1 for 1 hour. Cells were stimulated with PMA for 1 hour and labeled with 4-thiouridine (4sU) for another hour before collection for SLAM-seq. **H.** The distribution of log2 fold changes in nascent transcription following dTAGv1-induced TOP1 depletion relative to DMSO across all genes. The red dashed line indicates no change (log2 fold change = 0). **I.** The log2 fold change in nascent transcription following dTAGv1-induced TOP1 depletion relative to DMSO in NRSGs and RSGs. The red dashed line indicates no change (log2 fold change = 0). Statistical significance between groups is indicated (****P < 0.0001). **J.** Normalized expression levels of TCR signaling-responsive genes (RSGs) at 0 and 1 h after stimulation in DMSO- and dTAGv1-treated cells. Rows represent individual genes, and colors indicate relative expression levels. **K.** Schematic of the potential effects of negative supercoiling on highly transcribed genes and activated genes. **L.** Schematic of the experimental workflow. Cells were activated for 1 hour with PMA and subjected to DMSO or dTAGv1 for another hour, followed by 1 hour of 4-thiouridine (4sU) labeling before being collected for SLAM-seq. **M.** Schematic (upper) and boxplots (bottom) show expression levels of responsive genes (RSG, n= 295 genes) and non-responsive genes (NRSG, n= 2238 genes) at 0, 1, and 2 hours following PMA stimulation. **N.** The log2 fold change in nascent transcription following dTAGv1-induced TOP1 depletion relative to DMSO for responsive genes (RSG, n= 295 genes) and non-responsive genes (NRSG, n= 2238 genes) during the activation and highly transcribed phases. The red dashed line indicates no change (log2 fold change = 0), and blue dotted lines highlight the difference between the median responses of NRSGs across the two phases. Statistical significance between RSGs and NRSGs is indicated (***P < 0.001, ****P < 0.0001). **O.** CD69 expression in DMSO- and dTAGv1-treated cells following stimulation with increasing concentrations of PMA. Right: Quantification of CD69 geometric mean of fluorescence intensity (MFI) across PMA concentrations (n= 3 replicates).

We next investigated transcription-associated DNA supercoiling upon T-cell activation. Not surprisingly, genome-wide supercoiling profiling revealed that T-cell activation induced a global increase in transcription-associated supercoiling, with RSGs exhibiting a greater increase than NRSGs (Fig. 6E, F). Given our finding that accumulated supercoiling upon topoisomerase perturbation leads to transcriptional reduction, we tested whether the acute TOP1 depletion upon T-cell activation would influence gene activation. Indeed, we found that the global reduction in transcriptional activity following TOP1 depletion was more pronounced during T-cell activation than under steady-state conditions (Fig. 6G, H and Supplementary Fig. 8D). Importantly, RSGs exhibited a significantly greater reduction than NRSGs following acute TOP1 depletion (Fig. 6I– J). Together, these results indicate that genes undergoing rapid transcriptional activation accumulate more supercoiling for rapid topoisomerase resolution and thus are more sensitive to TOP1 depletion.

However, in addition to the fact that RSGs, but not NRSGs, undergo rapid transcriptional activation following T-cell stimulation, RSGs also exhibit higher transcriptional activity than NRSGs in general, which could contribute to their greater sensitivity to TOP1 depletion (Supplementary Fig. 8E). To distinguish these two effects and clarify whether the gene activation process itself imposes an additional sensitivity to supercoiling accumulation upon TOP1 depletion (Fig. 6K), we altered the timing of TOP1 depletion relative to T-cell activation. In the activation condition, TOP1 was depleted before T-cell stimulation (Fig. 6G), so that RSGs underwent transcriptional induction in the absence of TOP1. In the high-transcription condition, cells were first activated, and TOP1 was subsequently depleted (Fig. 6L), such that RSGs had already been induced and maintained a steady-state high transcriptional activity upon TOP1 depletion (Fig. 6M). This design allowed us to control for the general effect of TOP1 depletion on transcription due to different transcription levels and determine whether RSGs exhibit an additional sensitivity specifically during transcriptional activation. Notably, relative to the corresponding NRSGs, RSGs indeed showed a greater transcriptional reduction when TOP1 was depleted before their activation compared to when TOP1 was depleted after they had already reached a highly transcribed state (Fig. 6N). These results suggest that, in addition to negative supercoiling’s effect on steady-state transcriptional repression, accumulated negative supercoiling accumulation, if not efficiently resolved by topoisomerases, may further compromise rapid and robust transcription activation processes during cell-fate transitions.

In addition to measuring the transcriptional output via newly synthesized RNA, we also examined the expression of T-cell activation marker CD69 and observed a significant decrease upon TOP1 depletion, regardless of PMA and ionomycin concentrations applied for induction (Fig. 6O), confirming that the transcriptional defect ultimately translates into an attenuated cellular activation response. Together, these results support TOP1’s crucial function in resolving DNA topological tension that enables cells to accommodate rapid transcriptional upregulation during T-cell activation.

## Discussion

Our study reveals a regulatory function for transcription-generated DNA supercoiling that extends beyond its role as a byproduct and passive consequence of RNAPII elongation. By acutely depleting TOP1 and integrating measurements of DNA topology, nascent transcription, RNAPII dynamics, transcriptional bursting, and chromatin architecture, we find that transcription-generated negative supercoiling, if not resolved efficiently, represses transcriptional output at multiple levels. Negative supercoiling accumulation at the TSS is associated with increased promoter-proximal RNAPII pausing and slower RNAPII elongation, and TOP1 depletion also weakened enhancer-promoter interactions. It is the topoisomerase-mediated supercoiling relaxation that prevents transcription-generated negative supercoiling from accumulation, sustaining robust transcription in human cells.

A central implication of our findings is that the effect of negative supercoiling on transcription may be non-monotonic. Negative supercoiling has traditionally been considered favorable for transcription because underwinding DNA facilitates strand separation and promoter melting. Our results do not exclude the possibility that physiological level of negative supercoiling may facilitate promoter opening, but excessive accumulation of negative supercoiling following topoisomerase perturbation becomes inhibitory. This distinction may explain why genes with the highest transcriptional activity, and therefore those expected to generate the greatest transcription-associated topological stress, are most strongly inhibited by topoisomerase perturbation.

Our results also reveal an asymmetry between the two components of the twin-supercoiled domain. Although positive supercoiling ahead of RNAPII has generally been considered the major torsional impediment to elongation, changes in RNAPII elongation in our system are much more strongly associated with negative supercoiling at the TSS than with positive supercoiling at the TES. The behavior of genes in divergent, convergent, and tandem orientations further supports this conclusion. These findings suggest that negative supercoiling behind elongating RNAPII can impose a previously underappreciated constraint on transcription elongation across the human genome.

Together, our data suggest a unified feedback model. RNAPII generates negative supercoiling behind and positive supercoiling ahead as it transcribes DNA. Topoisomerase continuously relaxes transcription-associated supercoiling, preventing negative supercoiling from excessive accumulation. Upon the disruption of DNA topological homeostasis, negative supercoiling accumulates most strongly at highly transcribed loci. This altered topological state feeds back on the transcriptional system at several levels: chromatin interactions are weakened, promoter-proximal RNAPII pausing increases, and productive RNAPII elongation slows. These effects manifest at single-cell resolution as reductions in burst frequency and burst size and ultimately as lower transcriptional output and increased expression variability between individual cells.

Our experiments in T cells extend this model beyond steady-state transcription to the dynamic and physiologically relevant process of gene activation. To our knowledge, this is the first study to demonstrate a role for DNA topology in T-cell activation. We show that genes undergoing rapid transcriptional activation experience greater topological stress and are particularly sensitive to topological disruption such as TOP1 depletion, highlighting the importance of maintaining genome-wide DNA topological homeostasis to sustain rapid and robust T-cell activation. Given that T-cell activation is fundamental to T-cell development, differentiation, and effector functions^36^, our findings establish chromatin DNA topology as a previously unrecognized regulatory layer in T-cell biology. Further studies will be needed to define the broader roles of DNA supercoiling and DNA topoisomerase in T-cell development and differentiation and to determine whether dysregulated DNA topological homeostasis in T cells has a role in T-cell-mediated autoimmunity and antitumor immunity^37^.

In summary, our study establishes DNA supercoiling as an active component of chromatin organization and gene regulation in human cells. We propose that negative supercoiling accumulation upon topological homeostatic disruption is a transcriptional constraint. By controlling this balance, DNA topoisomerase coordinates chromatin looping and RNAPII dynamics in various steps of transcription regulation. This DNA-topology-mediated mechanism becomes particularly important during rapid gene upregulation and high-output transcription states such as T-cell activation.

## Limitations of the study

Although our genome-wide analyses upon supercoiling perturbation consistently associate negative supercoiling accumulation with increased RNAPII pausing, impaired elongation, and overall transcriptional repression, global rescue experiments and direct manipulation of DNA supercoiling at individual mammalian loci will be needed to fully establish the causality for supercoiling regulation of specific genes. The molecular mechanisms by which accumulated negative supercoiling impairs RNAPII dynamics also remain to be investigated, that may involve RNAPII backtracking, formation of alternative DNA structures and R-loops, and altered recruitment and activity of transcription, pause-release, and elongation factors.

## Methods

### Cell culture and G1 phase synchronization

HCT116 cells were cultured in McCoy’s 5A medium (Thermo Fisher Scientific, 16600108) supplemented with 10% FBS (fetal bovine serum, Sigma-Aldrich, F2442) and 1% PS (penicillin-streptomycin, Thermo Fisher Scientific, Cat#15140122) at 37 °C under 5% CO_2_ in the incubator. Jurkat T cells (E6-1 clone) were cultured at 37 °C with 5% CO2 in RPMI1640 medium containing 10% FBS and 1% PS. To synchronize the HCT116 cells, cells were cultured to reach 50%-70% confluence before adding lovastatin to 40 µM. Cells were washed with 10 mL McCoy’s 5A medium, then cultured with 5 mL medium for 4 hours after 24 hours of lovastatin treatment. To deplete TOP1 in HCT116-TOP1-mAID cells, the synchronized cells were incubated with 500 μM indole-3-acetic acid (IAA; Sigma, I3750-100MG-A) in the medium at 37°C for 2 hours. To inhibit TOP2 in HCT116 cells, the synchronized cells were incubated with 10 µM ICRF193 in the medium at 37 °C for 40 minutes. To deplete TOP1 in Jurkat-TOP1-dTAG cells, cells were incubated with 1 µM dTAGv1(MedChemExpress, Cat#: HY-145514C) in the medium at 37 °C for 1 hour. To activate Jurkat T cells, one million cells per mL were incubated with 0.5 × PMA/ionomycin (Thermo Fisher Scientific, Catalog number 00-4970-93) for the indicated duration.

### TTchem-seq and data analysis

TTchem-seq was performed following the published protocol with minor modifications^38^. Briefly, the synchronized cells were labeled with 500 μM 4sU (Thermo Fisher Scientific J60679-MD) for 20 minutes in the cell culture medium, before immediate quenching by removing the medium and adding QIAzol (Qiagen, cat. no. 79306). Total RNA was enriched using Chloroform (Thermo Fisher Scientific, cat. no. 43685), then purified using RNeasy Midi Kit (Qiagen, cat. no. 75144), and 400 ng of spike-in RNA from 4sU-labeled Drosophila S2 cells was added per 200 μg of total RNA. After RNA fragmentation to an average size of 250 bp using Covaris M220, nascent RNA labeled by 4sU was biotinylated by incubating with Biotium Biotin-XX MTSEA (Thermo Fisher Scientific, 50-196-4817) at room temperature in the dark for 30 min. The total RNA was then purified with Phenol/chloroform/isoamyl alcohol (25:24:1 (vol/vol); Thermo Fisher, cat. no.15593031), and the biotinylated nascent RNA was pulled down by μMACS streptavidin beads (Miltenyi Biotec, 130-074-101) and eluted with 100 mM DTT solution. The nascent RNA was then purified using the RNeasy MinElute Cleanup Kit (Qiagen, cat. no. 74204), followed by library preparation using the NEBNext rRNA depletion kit v2 (NEB, E7405) and NEBNext Ultra II directional RNA library prep kit (NEB, E7760S) before Illumina sequencing.

The raw data analysis follows the published protocol. Briefly, Paired-end reads were processed using fastp^39^ with automatic paired-end adapter detection (--detect_adapter_for_pe), poly-X trimming (--trim_poly_x), and a minimum retained read length of 25 nt (-l 25). Trimmed reads were aligned separately to the *Drosophila melanogaster* reference genome (BDGP6) and the human reference genome (GENCODE release 39) using STAR v2.7.10b^40^ in two-pass mode (-- twopassMode Basic), with gene-level counting and transcriptome alignments enabled (-- quantMode TranscriptomeSAM GeneCounts). Unmapped reads were excluded from BAM output (--outSAMunmapped None). Human-genome alignments were subsequently sorted with SAMtools^41^, and PCR duplicates were marked using Picard (Broad Institute) MarkDuplicates without removing duplicate reads (REMOVE_DUPLICATES=false).

Gene metaprofiles were generated using ngs.plot^42^, with normalization factors calculated based on the total number of reads mapped to the *Drosophila melanogaster* genome. Differentially expressed genes were identified using DESeq2^43^ as described in the protocol. Protein-coding genes were retained for differential expression analysis and all subsequent analyses involving fold changes and correlation analyses. The expression level of each gene was calculated by averaging across two biological replicates in the control cells.

To remove the effect of gene length on transcriptional fold changes, genes were grouped by length and the median log2 fold change (log2FC) and median gene length were calculated for each bin. A LOESS regression was fitted to the relationship between median log2FC and median gene length. The fitted model was then used to predict the gene-length-dependent component of log2FC for each individual gene. Length-normalized log2FC values were calculated by subtracting the predicted length-dependent effect from the observed log2FC, followed by an offset adjustment to preserve the overall center of the fold-change distribution.

### SLAM-seq and data analysis

The G1-synchronized and TOP1-depleted HCT116 cells were incubated with 500 µM 4sU for 1 hour before being harvested. The activated or TOP1-depleted Jurkat T cells were incubated with 500 µM 4sU for 1 hour before harvest. The total RNA was extracted following the manual of RNeasy Mini Kit (Qiagen). Next, RNA was subject to the SLAM-seq protocol with minor modifications. Briefly, 2 µg RNA was combined with 10 mM iodoacetamide (Sigma-Aldrich, St. Louis, MO, USA), 50 mM sodium phosphate buffer (pH 8.0), and 50% DMSO, followed by 15 min of incubation at 50°C. The reaction was stopped by adding 20 mM DTT, followed by RNA purification using 1.8X RNA beads (Aline Bioscience) and RNA elution with 11 µl nuclease-free water. Then, 500 ng RNA was subject to library construction using Quant-seq 3′ mRNA-Seq Library Prep Kit (Lexogen).

Data were first trimmed using cutadpt (v1.15) and then analyzed using SLAM-DUNK (v0.4.3)^44^ to map, filter, and quantify reads containing T-to-C mutations. Gene expression levels were calculated by averaging across biological replicates, with two replicates for HCT116-TOP1- mAID and control cells and three replicates each for Jurkat and Jurkat-TOP1-dTAG cells.

Differentially expressed genes were identified using DESeq2^43^, with the scaling factors calculated from the total RNA counts.

### ATMP-seq and data analysis

G1-synchronized HCT116 and Jurkat cells were treated with 90 μM and 180 μM Azide-TEG-TMP (ATMP; Berry & Associates, PS 5030), respectively, in growth medium for 10 min at 37°C. Cells were immediately placed on ice and irradiated with 360-nm UV-A light for 10 min at a total dose of 6 kJ/m², resulting in fewer than one monoadduct per 10 kb of DNA. Cells were washed with ice-cold PBS, and genomic DNA (gDNA) was purified using the DNeasy Blood & Tissue Kit (Qiagen) with RNase A treatment. Purified gDNA was subsequently sheared to an average fragment size of ∼600 bp using a Covaris M220 focused-ultrasonicator. For each 1 μg of gDNA, 1 μl of 10 mM DBCO-desthiobiotin was added, followed by incubation at 37 °C for 2 h with 950 r.p.m. interval shaking and purification using a Qiagen MinElute PCR purification kit. Biotin pulldown was performed with Dynabeads MyOne Streptevidin T1 beads, and the pulldown DNA was eluted with elution buffer (200 mM NaCl, 50 mM Tris pH 8.0 and 10 mM d-biotin) for 1 h at room temperature with rotation, further purified, and sonicated to 150 bp. ATMP libraries were amplified using the NEBNext Ultra II DNA library prep kit for Illumina sequencing. For the gDNA control, purified gDNA was first sonicated to an average fragment size of ∼600 bp, followed by ATMP incubation and UV-A irradiation under the same conditions used for the cellular samples to achieve a comparable ATMP adduct density. Sample and control DNA were subsequently processed in parallel using the same ATMP-seq enrichment and library-preparation workflow. Libraries were amplified using the same number of non-saturating PCR cycles and sequenced to comparable depths.

The raw reads of ATMP-seq were trimmed with Trimmomatic^45^ and aligned to the human reference genome (GENCODE release 39) with Bowtie 2^46^, sorted and indexed using SAMtools^41^, and deduplicated using Picard.

### ChIP-seq and data analysis

The ChIP-seq was performed as in the previous study, with minor modifications^47^. Briefly, G1-synchronized cells were crosslinked with 1% formaldehyde for 10 min at room temperature, and crosslinking was quenched with glycine for 5 min. Cells were washed twice with PBS, collected by centrifugation, snap-frozen, and stored at −80°C until further processing. For each ChIP, up to 20 × 10^6 cells were lysed in 0.5 mL ice-cold NP-40 lysis buffer for 5 min on ice, layered onto a sucrose cushion, and centrifuged at 13,000 × g for 10 min at 4°C to isolate nuclei. Nuclei were washed twice with PBS containing 1 mM EDTA and sequentially resuspended in glycerol buffer and nuclei lysis buffer. Following centrifugation, the chromatin pellet was washed twice with PBS containing 1 mM EDTA.

Chromatin was resuspended in 1 mL sonication buffer supplemented with 5 mM CaCl_₂_ and 600 U micrococcal nuclease (MNase) and digested for 15 min at 37°C with shaking at 700 rpm. MNase digestion was terminated by addition of EDTA and EGTA, and samples were placed on ice. Digested chromatin was further sheared using a Covaris M220 (PIP 75, 15% duty factor, 200 cycles/burst, 6 min) and clarified by centrifugation at 13,000 × g for 10 min at 4°C. Chromatin fragmentation was assessed using an aliquot of the lysate followed by DNA purification and TapeStation analysis.

Chromatin concentration was determined using the Bio-Rad DC Protein Assay. Clarified chromatin was divided into 300 µg aliquots and mixed with Spike-in (Active Motif, 53083) for each ChIP. Samples were incubated with 1 µg of Spike-in antibody (Active Motif, 61686) and 4 μg of RNA POLII (Active Motif, 39097) or Ser2P (Active Motif, 61083) antibody overnight at 4°C. Immune complexes were captured with 50 μL Protein A/G magnetic beads for 4–6 hours at 4°C. Beads were sequentially washed twice each with RIPA, RIPA containing 0.3 M NaCl, LiCl, and TE. Bound chromatin was reverse-crosslinked in 100 μL reverse-crosslinking buffer containing SDS and proteinase K by incubation overnight at 65°C with agitation. DNA was recovered, purified using a Zymo DNA purification kit, quantified, and used for library preparation with the NEBNext Ultra II DNA Library Prep Kit for Illumina.

ChIP-seq reads were adapter-trimmed using Trimmomatic^45^ and aligned to a combined hg38/dm6 reference genome using Bowtie2^46^. Alignments with MAPQ ≥10 were retained, and PCR duplicates were removed using Picard. Human and *Drosophila* reads were separated, and *Drosophila* spike-in read counts were used to calculate normalization factors. Spike-in-normalized bigWig tracks were generated using deepTools^48^ bamCoverage.

### NOTE-seq2 and data analysis

The cells were labeled with 250 µM 4sU for 2 hours, harvested by spinning at 300 x g for 3 minutes, resuspended in 500 µl (per million cells) ice-cold PBS, fixed by adding 500 µl (per million cells) 6% formaldehyde, and incubated on ice for 15 minutes. The cell fixation was quenched by adding 40 µl (per million cells) 2.5 M glycine and incubating on ice for 5 minutes. The cells were washed (hereafter, pellet cells at 500 x g for 5 minutes as needed) with SDP (0.5% SUPERase-In and 10 mM DTT in PBS), resuspended with SDP containing 0.2% Triton X-100, and incubated on ice for 5 minutes. Cells were pelleted and resuspended with 100 μl (per million cells) SDN buffer (0.5% SUPERase-In and 10mM DTT in nuclease-free water), followed by a quick mix with 600 µl 0.05N HCl (per million cells). After a 3-minute incubation on ice, cells were neutralized by adding 700 µl (per million cells) of 1 M Tris-HCl (pH 7.5), pelleted, and resuspended in 100 µl (per million cells) PS (0.5% SUPERase-In in PBS). When required, cells were supplemented with 10% DMSO and gradually frozen at −80°C for storage. For T-to-C conversion, frozen cells were thawed at 37°C and reacted with freshly prepared iodoacetamide (IAA). Specifically, 40 µl 100 mM fresh-made iodoacetamide (IAA, Millipore Sigma), 40 µl 500 mM Sodium phosphate buffer (To prepare Sodium phosphate buffer, mix 9.32 mL of 1 M Na_2_HPO_4_ and 0.68 mL of 1 M NaH_2_PO_4_ and add 10 mL of H_2_O. Adjust pH to 8.0.), and 20 µl nuclease-free water were sequentially added to the 100 µl cell suspension, before mixing with 200 µl DMSO. After 15-minute incubation at 50°C on a mixer, cells were put on ice with immediate addition of 8 µl 1 M DTT to quench the chemical conversion. Cells were then washed with 8.5 mL ice-cold PBS containing 0.2% Triton X-100. For DNA staining, cells were resuspended in 1 mL PBS containing 0.5% SUPERase-In and 2 μg/mL Hoechst 33342 and incubated for 10–15 min at room temperature protected from light. Cells were washed once with sorting buffer (0.5% BSA, 2 mM EDTA, 0.5% SUPERase-IN in PBS), passed through a 35–40 μm mesh, and resuspended in sorting buffer at approximately 2 × 10^6 cells per 300 μL. Samples were maintained on ice and protected from light until flow cytometric sorting into tubes containing 250 μL collection buffer (0.5% BSA, 0.5% SUPERase-IN in PBS). After sorting, cells were counted and loaded on the 10X Genomics platform, following the workflow according to the Chromium Next GEM-X Single Cell 3ʹ Reagent Kits v4 user guide (Rev A) with the following modifications: (1) After the GEM break (step 2.1a), the aqueous phase was transferred to a new PCR tube, with the addition of 1.5 µL 20%SDS and 5 µl 20 mg/ml protease K (Invitrogen), and then one-hour incubation at 53°C on a mixer. (2) After Dynabeads Cleanup (step2.1n), beads were resuspended with 10.5 µl MixtureA consisting of 50 mM Tris-HCl (pH 8.0), 60 mM NaCl, 5 mM MgCl_2_, and 1 mM dNTP. 10 µl elution was mixed with 10 µl MixtureB consisting of 2 mM GTP, 10% PEG8000, 16 mM DTT, 2U/µl Superase•In, 4 µM TSO (TSO is diluted from 1mM Template Switch Oligo from the kit), and 200 U Maxima H-minus, and incubated at 53°C for 45 minutes (second round of reverse transcription). (3) cDNA was amplified by mixing 50 µl Amp Mix, 15 µl cDNA primers, and 15 µl nuclease-free water, following the PCR program and cycles in the user guide (step2.2d). (4) The PCR products were purified with 80 µL PCRClean DX reagent, following step2.3. The downstream library preparation and sequencing were performed the same as the user manual.

Raw sequencing data were subjected to fastq calling, reads alignment, and quantification using Cell Ranger 9.0.1 with default parameters and standard pipeline. The gene expression matrix from the filtered_feature_bc_matrix was used to create a Seurat object using Seurat v4.5.0 with default parameters. Cellular barcodes (CBs) were retrieved from the Seurat object, and were used to subset the BAM file. Subsequently, confidently unique-aligned reads representing UMIs (corresponding to BAM tag: xf:i:25) were extracted from reads associated with the selected CBs. To quantify metabolically labeled and unlabeled transcripts, sites of T-to-C substitution with a base Phred Quality Score > 27 (A to G substitution in the reverse complementary strand) in each transcript were extracted and then filtered by background control without 4sU labeling and chemical conversion. The resultant transcripts containing T-to-C substitution were counted as newly synthesized transcripts and then corrected by a statistical model (detailed steps are available at https://github.com/lyuj2022/NOTE-seq).

### The barnyard experiment

The HCT116 cells and NIH3T3 cells were mixed at a 1:1 ratio, followed by the NOTE-seq workflow without T-to-C conversion. Reads were aligned to GRCh38_and_mm10-2020-A reference by cellranger 9.0.1 with cellular barcodes assigned to human, mouse, or multiplets. Briefly, cellular barcodes are initially classified as human or mouse by which genome has more total UMI counts for that barcode. Barcodes with total UMI counts that exceed the 10th percentile of the distributions for both human and mouse are called observed multiplets. Because cellranger only detects the observable multiplets (human, mouse), it computes an inferred multiplet rate by estimating the total number of multiplets, including (human, human) and (mouse, mouse). This is done by estimating via maximum likelihood the total number of multiplet GEMs from the observed multiplets and the inferred ratio of human to mouse cells.

### MNase-seq and data analysis

G1-synchronized cells were collected by centrifugation at 300 × g for 3 min at room temperature and crosslinked with 1% formaldehyde (FA) in PBS at a density of 2 × 10^6 cells/mL for 15 min at room temperature. Crosslinking was quenched by addition of 2.5 M glycine (50 μL/mL) for 5 min. Cells were aliquoted at 2 × 10^6 cells per sample, washed twice with PBS, and stored at −80°C. For MNase digestion, sample cells and spike-in cells (Drosophila S2) were combined at a 2:1 ratio and resuspended in 95 μL digestion buffer. Chromatin was digested with 1 KU MNase in a final volume of 100 μL for 15 min at 37°C, and the reaction was terminated by addition of 12.5 μL STOP buffer. Samples were treated with 3 μL RNase A (10 mg/mL) for 30 min at 42°C, followed by addition of 10 μL 10% SDS and 5 μL proteinase K (20 mg/mL) and incubation at 65°C overnight to reverse crosslinks. DNA was purified using a Zymo DNA purification kit. Purified DNA was resolved on a 2.5% agarose gel at 100 V for 40 min, and DNA corresponding to mononucleosomes (150–200 bp) was excised and recovered using the Zymo Gel DNA Recovery Kit. Recovered DNA was quantified using the Qubit High-Sensitivity assay, and sequencing libraries were prepared using the NEBNext Ultra II DNA Library Prep Kit.

MNase-seq reads were adapter-trimmed using Trimmomatic^45^ and aligned to a combined hg38/dm6 reference genome using Bowtie2^46^. PCR duplicates were removed, and properly paired reads with MAPQ ≥30 were retained. Human and *Drosophila* reads were separated for spike-in normalization, and mononucleosomal fragments (140–180 bp) were selected and processed using DANPOS^49^ for downstream nucleosome analysis.

### Micro-C and data analysis

Micro-C was performed following the protocol^50^. Briefly, G1-synchronized cells were initially crosslinked with 1% formaldehyde for 10 min at room temperature with rotation and quenched with 250 mM glycine for 5 min. After washing with cold PBS, cells were further crosslinked with 3 mM EGS for 45 min at room temperature, followed by quenching with 400 mM glycine for 5 min. Cell pellets were collected and stored at −80°C until further processing.

For each batch of cells, the optimal micrococcal nuclease (MNase) concentration was determined by titration. Cells were permeabilized in Micro-C buffer (10 mM Tris-HCl, pH 7.5, 50 mM NaCl, 5 mM MgCl_₂_, 1 mM CaCl_₂_, 0.2% NP-40, and protease inhibitors) and digested with serially diluted MNase for 15 min at 37°C. Digestion conditions were selected based on agarose-gel analysis to obtain a prominent mononucleosome band at approximately 150 bp and a dinucleosome band at approximately 320 bp, with minimal subnucleosomal DNA. Preparative samples were then digested under the selected MNase condition for 15 min at 37°C with shaking at 800 rpm, and the reaction was terminated with EGTA. An aliquot was retained as an input control to assess MNase digestion.

MNase-digested chromatin was subjected to DNA end processing with T4 polynucleotide kinase and Klenow fragment, followed by incorporation of biotin-dATP and biotin-dCTP and proximity ligation with T4 DNA ligase. The reaction was subsequently treated with Exonuclease III, followed by proteinase K/SDS treatment and overnight incubation at 65°C. DNA was purified and resolved on a 1.5% agarose gel, and the proximity-ligated dinucleosomal DNA fraction was excised and recovered using a gel DNA recovery kit. Purified DNA was quantified using the Qubit dsDNA High-Sensitivity assay.

Biotinylated ligation products were enriched using streptavidin beads and subjected to end repair/A-tailing, adaptor ligation, and USER enzyme treatment. The number of library-amplification cycles was empirically determined by a preliminary PCR to avoid overamplification, and the final libraries were amplified using high-fidelity DNA polymerase. Amplified libraries were purified using SPRIselect or NEBNext sample-purification beads, and library fragment-size distributions were assessed using a TapeStation before sequencing.

Micro-C libraries were processed from raw valid pairs files for DMSO (control) and IAA (auxin-induced TOP1-AID degradation) conditions in HCT116-TOP1-mAID cells. Contact matrices were generated using cooler^51^ cload pairix for different resolutions followed by matrix balancing (*cooler balance*, ICE normalization) on the original valid pairs files.

#### Loops calling

Chromatin loops were called independently for DMSO and IAA at 4 kb resolution using Mustache (v1.3.3)^52^ with sparsity threshold -st 0.2 and p-value threshold -pt 0.1. All loops were filtered to FDR < 0.05 prior to downstream comparison to call high-confident loops. These loops were classified into five mutually exclusive categories based on anchor-level comparison between conditions, using exact bin-coordinate matching: **Shared** (both anchors matched exactly between DMSO and IAA), **Shrunk** (one anchor shared; the non-shared anchor moved closer in IAA, decreasing loop span), **Expanded** (one anchor shared; the non-shared anchor moved farther in IAA, increasing loop span), **Lost** (a DMSO loop anchor with no corresponding match in IAA), **Gained** (an IAA loop anchor with no corresponding match in DMSO). Loops matching was performed using a custom Python script where each DMSO loop searched for candidate IAA loops sharing at least one anchor. Loop span was defined as the genomic distance between anchor midpoints. IAA loops sharing no anchor with any DMSO loop were classified as Gained. For each category, loop span (anchor-midpoint distance), FDR, and chromosomal distribution were characterized separately at 4 kb. Distributions were compared using the Mann-Whitney U test for all pairwise category comparisons, with Benjamini-Hochberg correction applied within each (resolution, property) family of ten pairwise tests. Promoters were defined as GENCODE v44 protein-coding transcript TSS positions ± 2 kb. Enhancers were defined as ENCODE candidate cis-regulatory elements (cCREs, GRCh38). A loop was classified as an E-P loop if one anchor overlapped a promoter and the other anchor overlapped an enhancer (either orientation). E-P loop percentages were calculated for each category.

#### Stripes calling

Architectural stripes were called independently for DMSO and IAA at 4 kb resolution using Stripenn (v1.3.3)^53^. Raw Stripenn output was filtered to Stripiness > 0 (Stripenn’s internal call-quality metric, reflecting gradient contrast and signal continuity in the observed/expected-normalized contact matrix). Stripe lengths were characterized per sample. These stripes were compared between conditions by base-region (fixed anchor) overlap: a DMSO stripe was considered matched to an IAA stripe if their base regions overlapped, with the best match (largest overlap span) selected in cases of multiple candidates. Matched stripes were classified as Shared (base overlap present); unmatched DMSO stripes were classified as Lost, and unmatched IAA stripes as Gained. To assess whether stripe weakening reflected genuine biological change versus a quality-threshold artifact, DMSO stripes (QC-filtered, Stripiness > 0) were matched against IAA’s raw counterparts, unfiltered Stripenn output using anchor-proximity matching (20 kb tolerance), so that “disappeared” reflected genuine absence from Stripenn’s raw calls rather than exclusion by the quality filter. The two-sample Kolmogorov-Smirnov test was used to compare the shape of the DMSO and IAA (surviving-stripe) strength distributions.

#### Insulation and Topologically Associating Domains (TADs)

Diamond-window insulation scores were computed from 1 kb resolution cool matrices using cooltools^54^.insulation, with window sizes of 50 kb, and 100 kb, ignoring the first two diagonals, using ICE-balanced weights, and an adaptive (Li) threshold for boundary calling. TADs were defined as the genomic intervals between consecutive passing boundaries within each chromosome. TADs we filtered by excluding TADs shorter than 3× the diamond window size.

#### A/B Compartments

Compartment analysis was performed at 100 kb and 250 kb cool resolutions. The first eigenvector (E1) of the observed/expected cis-contact matrix was computed using *cooltools.eigs.cis*, using ICE-balanced weights. Eigenvector sign was oriented using genome-wide GC content (computed from the hg38 reference sequence via bioframe^55^) as the phasing track, such that positive E1 corresponds to the GC-rich, gene-dense A compartment by convention. Bins were assigned to compartment A (E1 > 0) or B (E1 ≤ 0); bins with undefined E1 (e.g., low-coverage or unmappable regions) were left unassigned. Compartment switching between DMSO and IAA was assessed per bin by comparing compartment calls at matched genomic positions. To visualize compartment organization directly, E1 profiles were plotted as filled area tracks for DMSO and IAA, with positive E1 (A compartment) filled above zero and negative E1 (B compartment) filled below zero, at 100 kb resolution. Tracks were generated for two representative 70 Mb regions (chr11:60–130 Mb and chr4:40–110 Mb) to allow direct visual comparison of compartment domain structure between conditions.

#### Genome-Wide Contact Scaling (P(s) Curves)

Genome-wide cis contact probability as a function of genomic distance was computed using cooltools.expected_cis at 4 kb. Per-chromosome expected values were aggregated into a genome-wide curve weighted by the number of valid pixels at each genomic distance. The local slope of the log-log P(s) curve was computed via numerical differentiation to identify the loop-extrusion-associated shoulder distance in each condition.

#### Supercoiling Domain Analysis

To test whether loop disruption under TOP1 depletion is associated with DNA supercoiling state, genome-wide supercoiling domains were defined from ATMP-supercoiling sequencing tracks (BigWig, 500 kb bins; HCT116-TOP1-mAID, G1-synchronized, auxin-treated). Bins with signal greater than zero were classified as negative-supercoiling domains, and bins with signal less than zero as positive-supercoiling domains. Mustache loop calls for DMSO and IAA (4 kb resolution) were filtered to FDR < 0.05 prior to classification. For each loop, both anchors were independently assigned a supercoiling domain class based on the domain bin overlapping the anchor midpoint. A loop was classified as a “negative” or “positive” supercoiling loop if both anchors fell within the same domain type. The number of loops in each supercoiling category was compared between DMSO and IAA.

### TOP1-dTAG Jurkat cell construction

For generation of the TOP1-dTAG donor plasmid (Hygromycin), the mAC-POLR2A donor plasmid was used as the parental backbone. The vector backbone was PCR-amplified from the parental plasmid, and two TOP1 genomic DNA fragments flanking the intended N-terminal tagging site were amplified from genomic DNA to serve as homology arms. A fragment containing the tagging/selection cassette was amplified from the mAC-POLR2A donor, and the four PCR products were assembled by Gibson assembly to generate the mAC-TOP1 donor plasmid. The mAID-based tagging region was then modified by sequence replacement to introduce the FKBP12^F36V^-based dTAG module, yielding the FKBP-V-AC-TOP1_N donor plasmid. The resulting construct was designed for homology-directed insertion of the degron tag at the N-terminus of endogenous TOP1. For generation of TOP1-dTAG donor plasmid (Blasticidin). The pMK348 (BSD-P2A-mAID-mClover) was used as the parental backbone. TOP1 genomic fragments encompassing sequences upstream and downstream of the N-terminal sgRNA target site were introduced into the parental vector to generate the TOP1 homology arms for homology-directed repair. The mAID degron sequence was subsequently replaced with FKBP12^F36V^, while retaining the BSD-P2A selection cassette and mClover fluorescent reporter. The resulting pMK348 (BSD-P2A-FKBP12^F36V^-mClover-TOP1_N) donor plasmid contained the TOP1 homology arms flanking the BSD-P2A-FKBP12^F36V^-mClover tagging cassette. The TOP1-targeting Cas9–gRNA plasmid was constructed using the pX330-U6-Chimeric_BB-CBh-hSpCas9 backbone. Complementary oligonucleotides encoding the 20-nt TOP1_N sgRNA sequence (5′-GCTCCGGGCCGACATGAGTG-3′) were annealed and cloned into the sgRNA expression cassette of pX330.

To construct the TOP1-dTAG cell line, 1 × 10^6 cells were collected by centrifugation at 90 × g for 10 min at room temperature and resuspended in 100 μL supplemented SE Cell Line Nucleofector Solution (82 μL SE solution and 18 μL Supplement 1). Cells were mixed with 1 μg TOP1 donor plasmid and 1 μg TOP1 Cas9–gRNA plasmid (Hygromycin and Blasticidin) and nucleofected using the CL-120 program on the Nucleofector X Unit. Immediately after nucleofection, 300 μL prewarmed RPMI 1640 recovery medium was added, and cells were allowed to recover for 10 min at 37°C before transfer to prewarmed culture medium. Following 48 h of recovery, cells were subjected to selection with 500 μg/mL hygromycin B and 10 µg/µL for 4 weeks. Following antibiotic selection, cells were diluted to an average density of 0.5 cells per well and distributed into two 96-well plates for single-cell cloning. Clones were expanded for approximately 4 weeks before further validation.

### Flow cytometry

Cell-cycle distribution was assessed by propidium iodide (PI) staining followed by flow cytometry. Briefly, Cells were detached with 0.5 mL trypsin for 2 min at 37°C, neutralized with 1 mL culture medium, and 2.5 × 10^6 cells were collected by centrifugation at 250 × g for 5 min. Cells were washed once with PBS, resuspended in 0.5 mL PBS, and fixed by dropwise addition of 2 mL methanol with continuous vortexing, followed by incubation on ice for 1 h. An aliquot (1 mL) of the fixed-cell suspension was collected by centrifugation at 500 × g for 5 min and resuspended in 500 μL staining buffer containing 10 μg/mL RNase A (Thermo Scientific) and 20 μg/mL PI (Thermo Scientific). Cells were incubated for 10 min at room temperature, protected from light, then passed through a 40-μm cell strainer and kept on ice until flow-cytometric analysis of DNA content and cell-cycle distribution.

TOP1 depletion was assessed by monitoring mClover fluorescence using flow cytometry. Jurkat-TOP1-FKBP cells were treated with 1 μM dTAGv1 at 37°C for the indicated durations and subsequently harvested for flow-cytometric analysis.

To assess T cell activation, Jurkat cells (1.25 × 10^5 per well) were seeded in 48-well plates containing 450 μL culture medium and treated with 1 μM dTAGv1 or an equivalent volume of DMSO for 1 h. Cells were subsequently stimulated with PMA/ionomycin at the indicated concentrations (0, 0.1×, 0.5×, or 1×) for an additional 4 h. Cells were harvested by centrifugation at 300 × g for 5 min and stained in 50 μL cell-staining buffer containing APC-conjugated anti-human CD69 antibody for 30 min at 4°C, protected from light. Cells were washed once with 1 mL cold PBS, resuspended in 300 μL cold PBS, and passed through a 40-μm cell strainer. Samples were maintained on ice and protected from light until flow-cytometric analysis. Cells were analyzed by BD LSRFortessa SORPI, and data were analyzed by FlowJo v10.

### Western blot

Following TOP1 depletion by IAA or dTAGv1 incubation. Cells were lysed in RIPA buffer supplemented with protease inhibitors for protein extraction. The cell lysate was quantified by DC protein assay (BioRad), followed by 95°C protein denaturation in 1x SDS loading buffer. Equal amounts of protein samples were loaded to and separated by 10% SDS-PAGE and then transferred onto a PVDF membrane (Millipore). The membrane was blocked in 5% non-fat milk dissolved in Tris-buffered saline with Tween-20 (TBST) for 1 hour and then washed with TBST three times. The membrane was then incubated with rabbit anti-mouse primary antibodies against TOP1(Abcam, ab109374) or beta-actin (Abcam, ab272085) overnight at 4°C, and with horseradish peroxidase-labeled secondary antibody (Abcam, ab97080) at room temperature for 1 hour. The protein signals were detected by ECL Kit (Thermo Scientific, 34580) on Cytiva800.

### Sequencing

All libraries were sequenced on the AVITI sequencing Platform (AmpSeq, LLC). For TTchem-seq, 2 x 150 cycles paired-end sequencing was performed. For ATMP-seq, ChIP-seq, MNase-seq, and Micro-C, 2 x 75 cycles paired-end sequencing was performed. For SLAM-seq, 150-cycle single-end sequencing was performed. For NOTE-seq2, sequencing was performed according to the parameters specified in the Chromium Next GEM Single Cell 3′ Kit v4 User Guide, except that Read 2 was extended to 120 cycles.

## Acknowledgements

We thank Charan Vemuri and Qian Yao from the Chen lab for their help in ChIP-seq and ATMP-sea data analysis, respectively. We thank previous lab member Linyin Zhu for sharing the spike-ins for TTchem-seq. The HCT116-TOP1-mAID cell line is a gift from L. Baranello (Karolinska Institute). The Jurkat T cell line is a gift from Peter D. Aplan (National Cancer Institute). This study used the Biowulf Linux cluster at the National Institutes of Health. This work was supported by the Intramural Research Program of the National Institutes of Health, National Cancer Institute.

## Author contributions

J.L. conceived the project, performed most of the experiments, analyzed most of the data, prepared figures, and wrote the manuscript. A.A. performed Micro-C and analyzed the data. I.B. performed ATMP-seq. S.L. constructed the TOP1-dTAG plasmids and provided helpful suggestions. C.C. conceived and supervised the project and edited the manuscript.

---

**Supplementary Figure 1.**
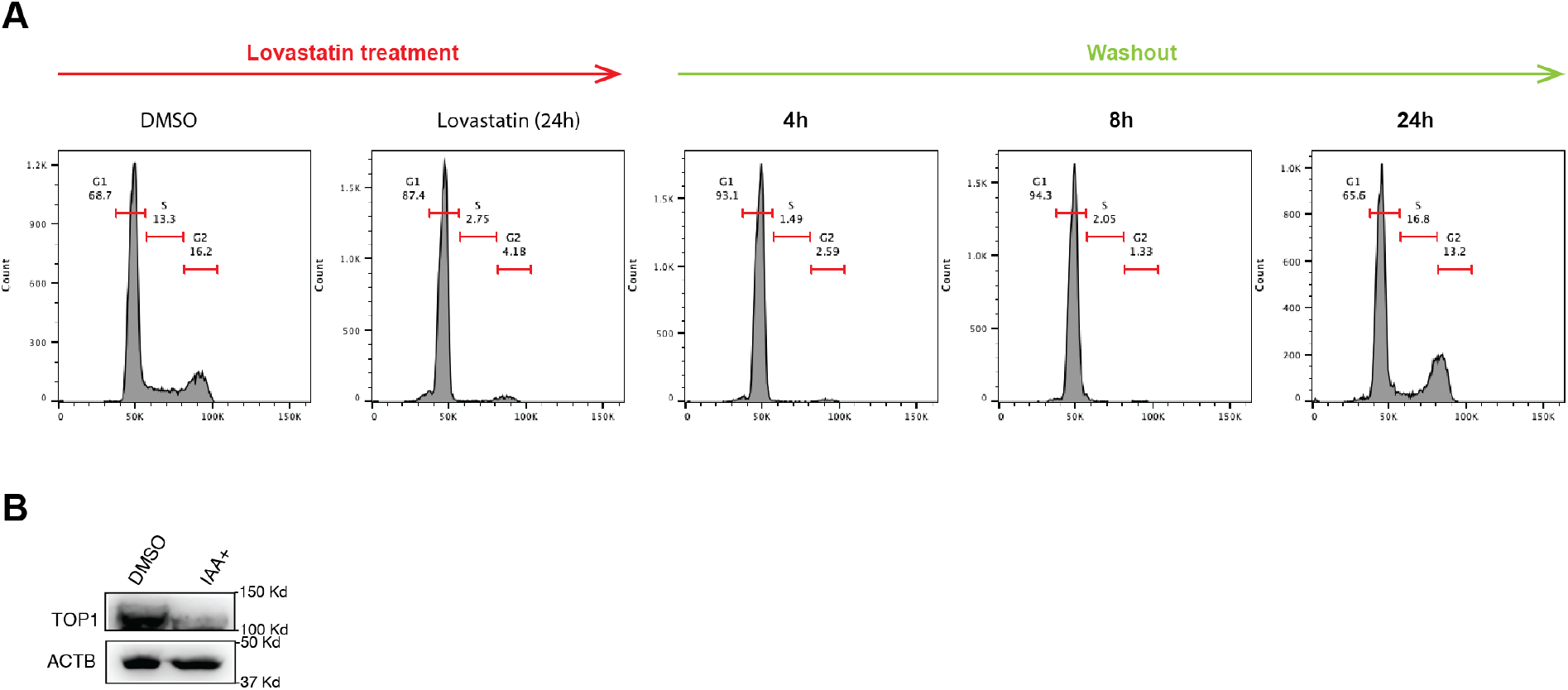
G1 synchronization and acute TOP1 depletion. **A.** Cell-cycle distributions in DMSO-treated cells, after 24 h of lovastatin treatment, and at 4, 8, and 24 h following lovastatin washout. The percentages of cells in G1, S, and G2 phases are indicated. **B.** Acute TOP1 depletion after 2 hours of IAA treatment.

**Supplementary Figure 2.**
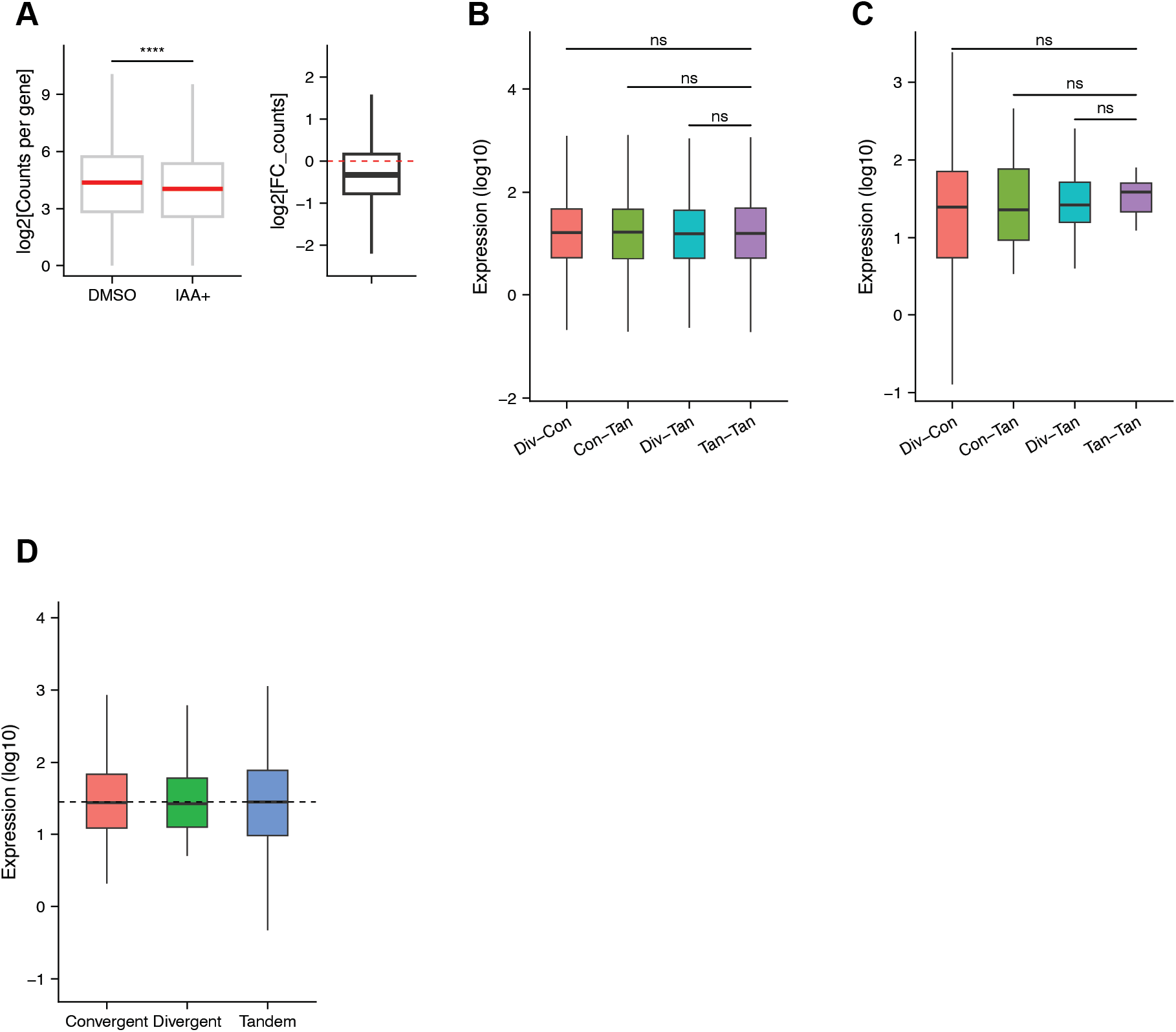
Negative supercoiling accumulation at the TSS represses transcriptional activity. **A.** Left: Transcription activity of genes upon TOP1 depletion (IAA) versus control (DMSO), measured by SLAM-seq. Right: Fold change in transcription activity of genes upon TOP1 depletion. The transcription level is the average of two replicates (n= 10435 genes). **B-D.** Expression activity of genes across different gene contexts. No significant differences in gene expression were observed among the indicated groups (ns, not significant).

**Supplementary Figure 3.**
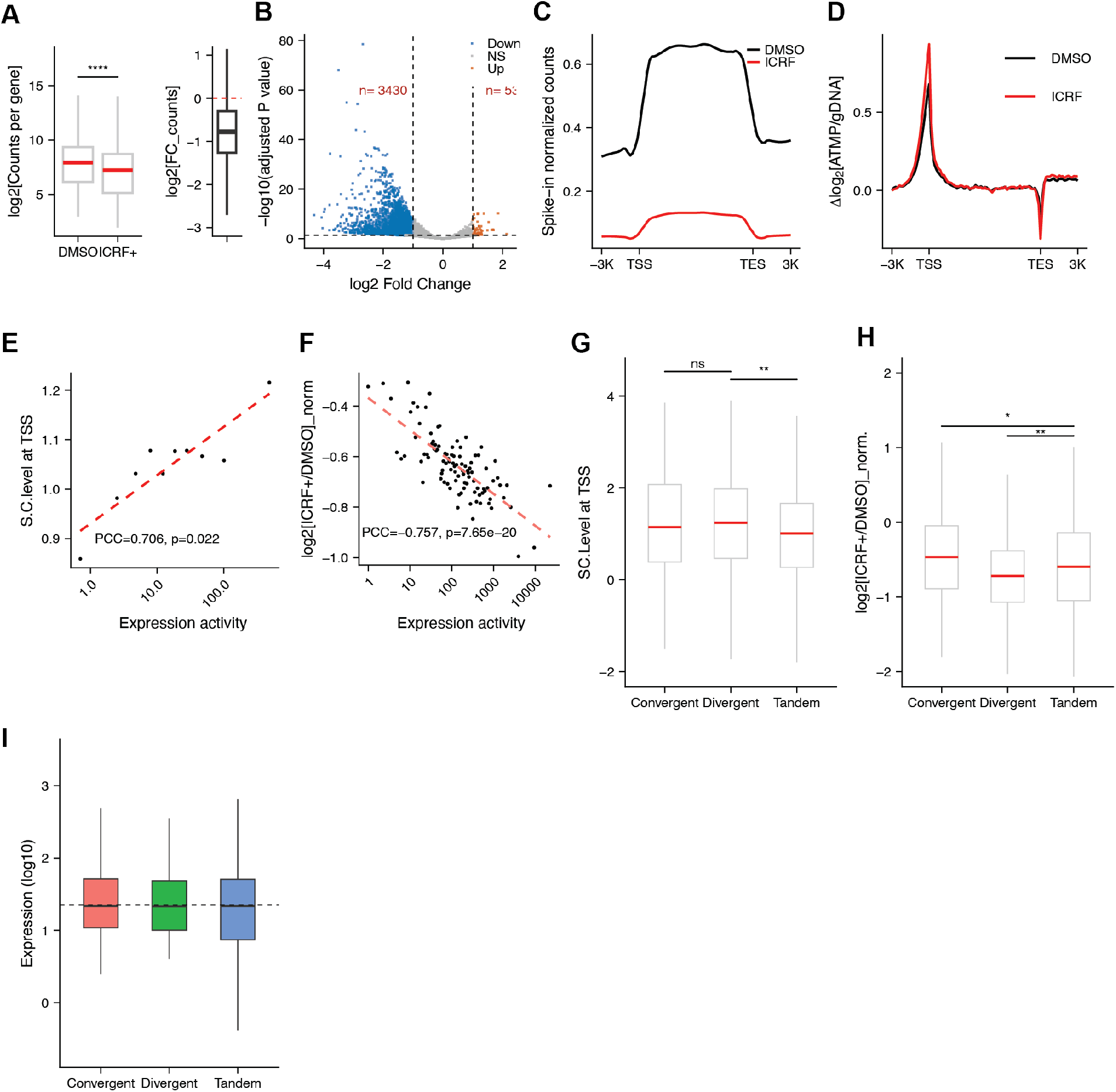
Negative supercoiling accumulation at the TSS by TOP2 perturbation represses transcription. **A.** Left: Transcription activity of genes upon TOP1 depletion (IAA) versus control (DMSO), measured by TTchem-seq. Right: Fold change in transcription activity of genes upon TOP1 depletion. n= 16333 genes. **B.** Genes with significantly changed transcription activity (p-adj ≤ 0.05, |log2(fold change)| ≥ 1, n= 15699 genes). **C.** Nascent transcripts along the gene body of genes upon ICRF193 versus DMSO treatment. **D.** Supercoiling level around genes upon ICRF193 or DMSO treatment, relative to the supercoiling level at the gene body (n= 19,804 genes). **E.** The correlation between S.C. level at the TSS and transcriptional activity of genes in the DMSO condition. Genes were grouped into bins of 1,000 genes, with each dot representing the median of one bin. The dashed red line indicates the linear regression fit (n= 10 bins for 9314 genes). **F.** The correlation between transcriptional activity of genes in DMSO condition and the normalized fold change in transcriptional activity of genes upon ICRF193 treatment relative to DMSO control. Genes were grouped into bins of 100 genes, with each dot representing the median of one bin. The dashed red line indicates the linear regression fit (n= 100 bins for 9942 genes). **G.** S.C. level at the TSS of genes across different gene contexts. The distance between the focal gene and its flanking genes is greater than 4,500 bp on one side and less than 1,500 bp on the other. *n* = 271, 464, and 400 for convergent, divergent, and tandem gene contexts, respectively. **H.** Normalized fold change in transcriptional activity of genes in (G), following ICRF193 treatment relative to DMSO across different gene contexts. **I.** Expression activity of genes in (G) across different gene contexts. No significant differences in gene expression were observed among the indicated groups. In (G) and (H), red horizontal lines indicate group medians, and the dashed line indicates the median of the group in the tandem context. Pearson correlation coefficient (PCC) and associated *P* value are shown as indicated in (E) and (F). Statistical significance is indicated in (A), (G), and (H) (*P < 0.05, **P < 0.01, ****P < 0.0001; ns, not significant).

**Supplementary Figure 4.**
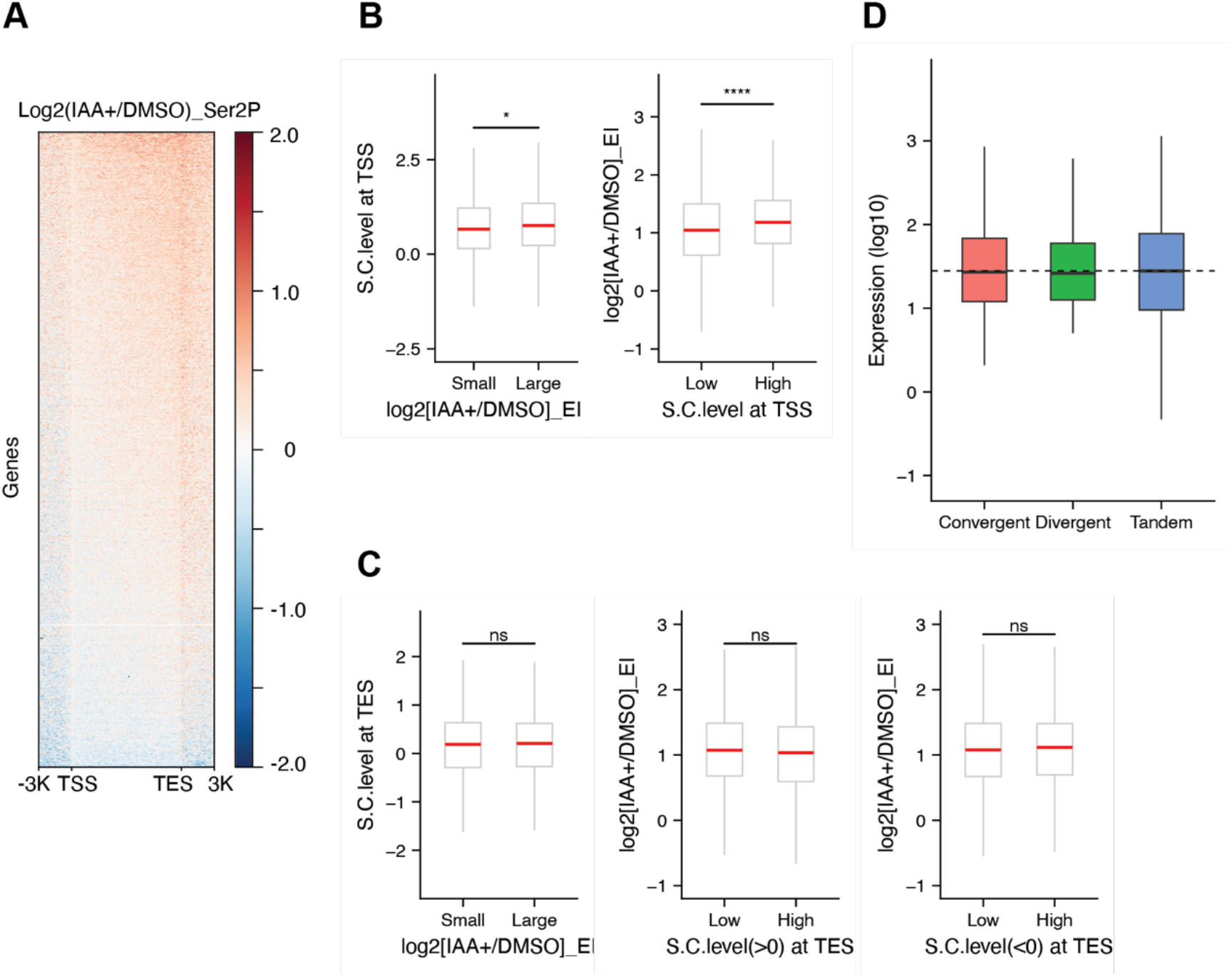
Supercoiling accumulation at the TSS slows down RNAPII elongation. **A.** The log2 fold change in RNAPII Ser2P ChIP-seq signal following IAA-induced TOP1 depletion relative to DMSO across genes, spanning 3 kb upstream of the transcription start site (TSS), the gene body from TSS to transcription end site (TES), and 3 kb downstream of the TES. Red and blue indicate increased and decreased Ser2P signal, respectively. **B.** Left: Comparison of S.C. level at the TSS in control between genes showing the smallest and largest changes in elongation indices upon TOP1 depletion relative to the DMSO control. Right: Comparison of fold change in elongation indices between genes with the lowest and highest supercoiling levels in the control. **C.** Left: Comparison of S.C. level in control at the TES between genes showing the smallest and largest changes in elongation indices upon TOP1 depletion relative to DMSO control. Middle and Right: Comparison of fold change in elongation indices between genes with the lowest and highest supercoiling levels in control. In B and C, each category includes 1000 genes. Red lines indicate medians. **D.** Expression activity of genes across different gene contexts. No significant differences in gene expression were observed among the indicated groups. Statistical significance is indicated in B and C (*P < 0.05, ****P < 0.0001; ns, not significant).

**Supplementary Figure 5.**
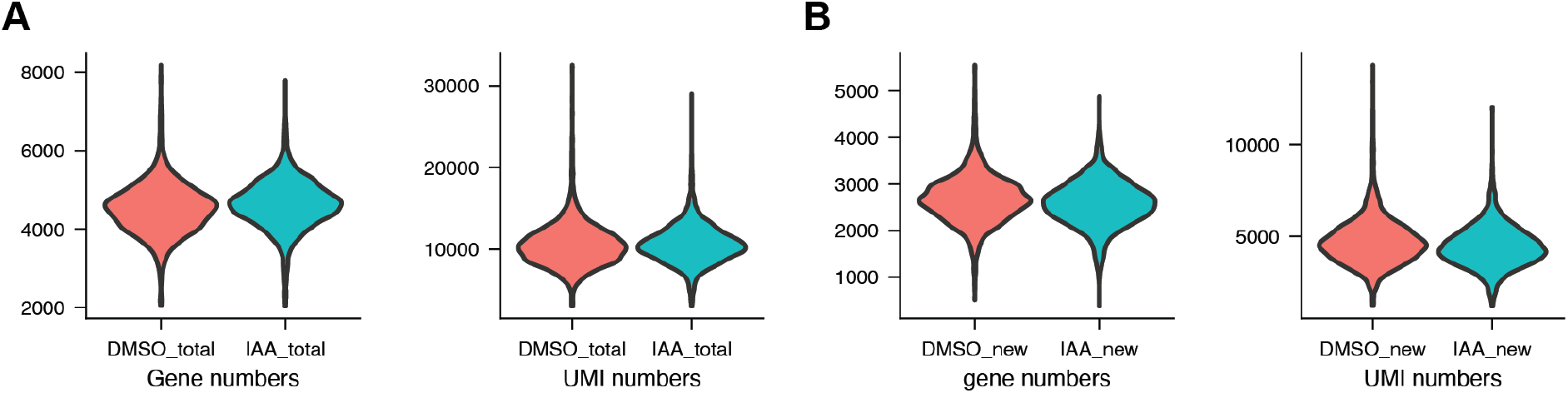
Genes and transcripts detected by NOTE-seq2. **A-B.** The distributions of detected gene numbers and UMI counts for **(A)** total RNA and **(B)** newly synthesized RNA in DMSO- (n=3220 cells) and IAA- (n=3160 cells) treated cells.

**Supplementary Figure 6.**
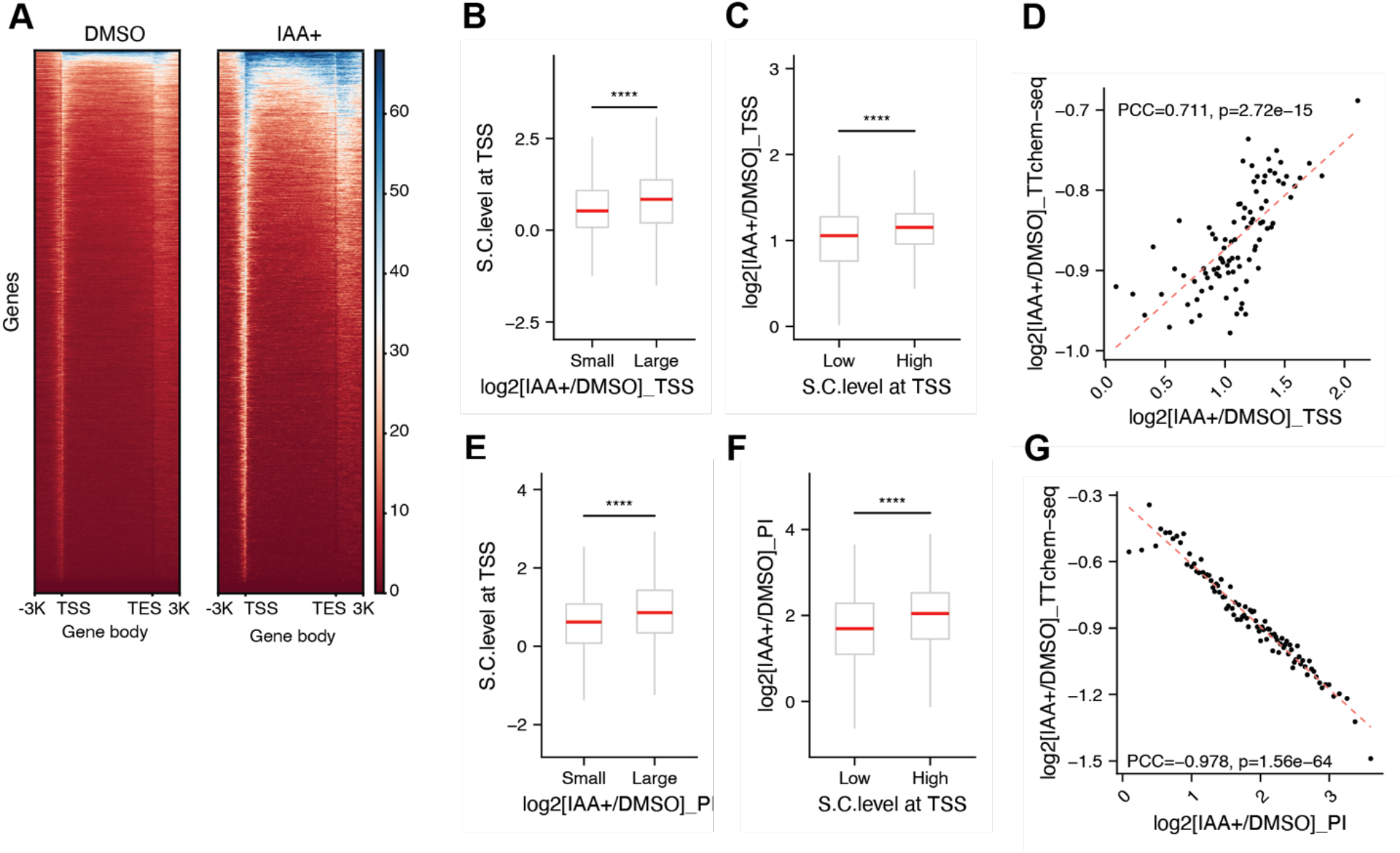
Negative supercoiling at the TSS increases RNAPII occupancy and pausing at promoter regions. **A.** RNAPII ChIP-seq signal across genes in DMSO- and IAA-treated cells, spanning 3 kb upstream of the transcription start site (TSS), the gene body from TSS to transcription end site (TES), and 3 kb downstream of the TES. **B and E.** Comparison of S.C. level at the TSS in the DMSO control between genes showing the smallest and largest changes in RNAPII ChIP-seq signal at the TSS (B) or in the pausing index (E) upon TOP1 depletion relative to the DMSO control. **C and F.** Comparison of fold change in RNAPII ChIP-seq signal at the TSS (C) or in the pausing indices (F) between genes with the lowest and highest supercoiling levels at the TSS in the DMSO control. In B, C, E, and F, each category includes 1000 genes. Red lines indicate medians. Statistical significance is indicated (****P < 0.0001). **D and G.** Correlation between fold change in transcription activity and the fold change in RNAPII binding at the TSS (D) or in pausing indices (G) of genes upon TOP1 depletion relative to the DMSO control. Genes were grouped into bins of 1,00 genes, with each dot representing the median of one bin. The dashed red line indicates the linear regression fit.

**Supplementary Figure 7.**
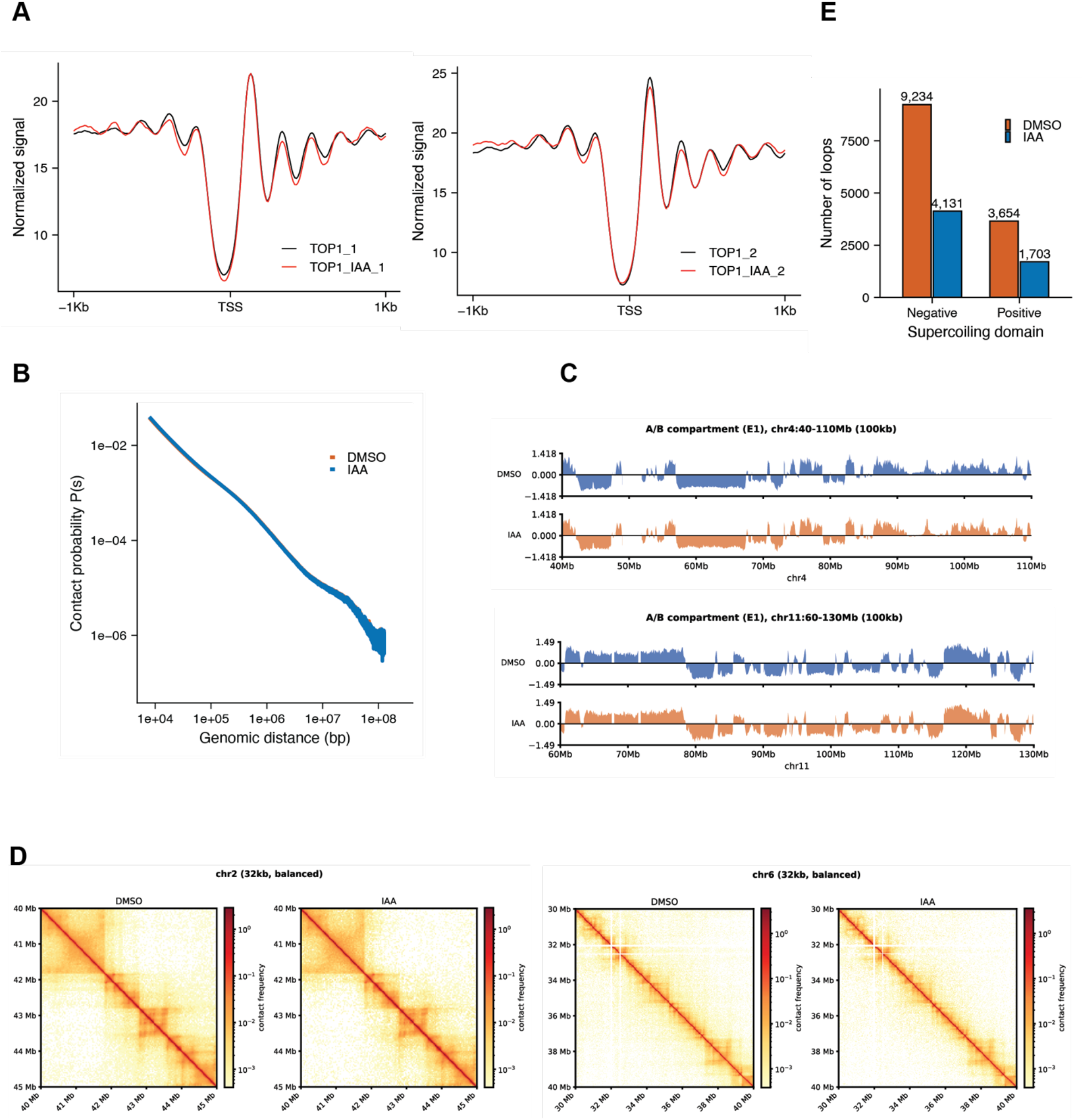
Nucleosome occupancy and large-scale 3D genome organization remain unchanged upon TOP1 depletion. **A.** Normalized MNase-seq signals centered at transcription start sites (TSSs), spanning 1 kb upstream and downstream, in DMSO- and IAA-treated cells for two replicates. **B.** Genome-wide contact probability as a function of genomic distance in DMSO- and IAA-treated cells. **C.** Representative eigenvector profiles show A/B compartment organization at 100-kb resolution across regions of chromosomes 4 and 11 in DMSO- and IAA-treated cells. **D.** Hi-C contact maps at 32-kb resolution show representative regions of chromosomes 2 and 6 in DMSO- and IAA-treated cells. Color intensity indicates contact frequency. **E.** The number of chromatin loops located within negative and positive supercoiling domains in DMSO- and IAA-treated cells. The number of loops is indicated above each bar.

**Supplementary Figure 8.**
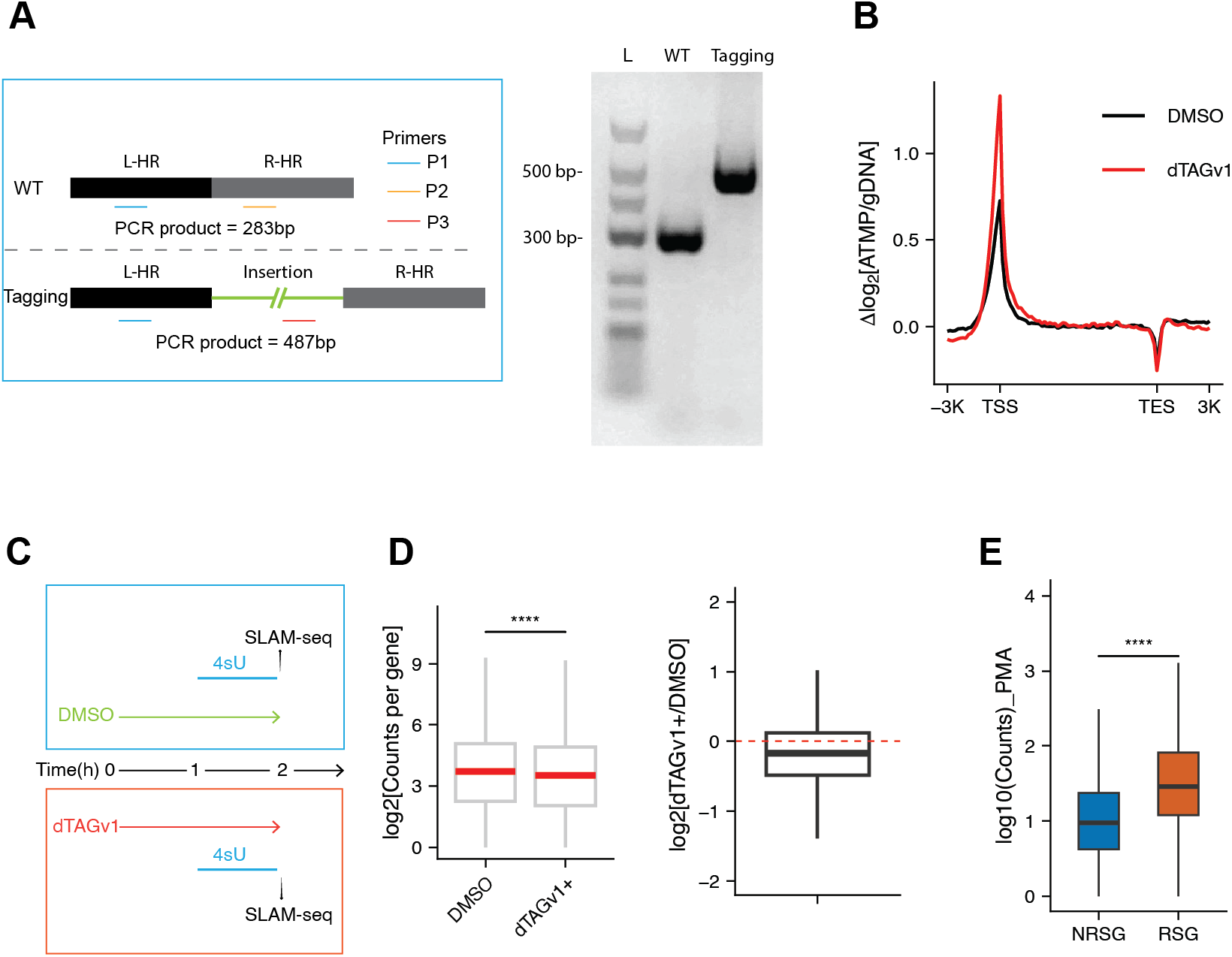
Negative supercoiling accumulation upon TOP1 depletion compromises transcriptional activity and rapid gene activation in T lymphocytes. **A.** Validation of targeted genomic tagging by PCR. Left, schematic of the PCR strategy used to distinguish wild-type (WT) and tagged alleles, with expected PCR products of 283 bp and 487 bp, respectively. Primer positions (P1–P3), left and right homology regions (L-HR and R-HR), and the inserted sequence are indicated. Right, agarose gel electrophoresis showing PCR products of the expected sizes for WT and tagged alleles. L, DNA ladder. **B.** Supercoiling level around genes upon dTAGv1or DMSO treatment, relative to the supercoiling level at the gene body (n= 19872 genes). **C.** Schematic of the experimental workflow. Cells were treated with DMSO or dTAGv1 for 1 hour and labeled with 4-thiouridine (4sU) during the final hour before collection for SLAM-seq. **D.** Left, boxplots show log2-transformed nascent RNA counts per gene in DMSO- and dTAGv1-treated cells. Right, distribution of log2 fold changes in nascent transcription following dTAGv1-induced TOP1 depletion relative to DMSO. The red dashed line indicates no change (log2 fold change = 0). **E.** The log10-transformed nascent RNA counts following PMA stimulation in TOP1-depleted cells for non-responsive genes (NRSGs, n= 3779) and responsive genes (RSGs, n= 541). In D and E, Statistical significance is indicated (****P < 0.0001).

